# On the path to reference genomes for all biodiversity: lessons learned and laboratory protocols created in the Sanger Tree of Life core laboratory over the first 2000 species

**DOI:** 10.1101/2025.04.11.648334

**Authors:** Caroline Howard, Amy Denton, Benjamin Jackson, Adam Bates, Jessie Jay, Halyna Yatsenko, Priyanka Sethu Raman, Abitha Thomas, Graeme Oatley, Raquel Vionette do Amaral, Zeynep Ene Göktan, Juan Pablo Narváez Gómez, Isabelle Clayton Lucey, Elizabeth Sinclair, Michael A. Quail, Mark Blaxter, Kerstin Howe, Mara K. N. Lawniczak

## Abstract

Since its inception in 2019, the Tree of Life programme at the Wellcome Sanger Institute has released high-quality, chromosomally-resolved reference genome assemblies for over 2000 species. Tree of Life has at its core multiple teams, each of which are responsible for key components of the ‘genome engine’. One of these teams is the Tree of Life core laboratory, which is responsible for processing tissues across a wide range of species into high quality, high molecular weight DNA and intact RNA, and preparing tissues for Hi-C. Here, we detail the different workflows we have developed to successfully process a wide variety of species, covering plants, fungi, chordates, protists, arthropods, meiofauna and other metazoa. We summarise our success rates and describe how to best apply and combine the suite of current protocols, which are all publicly available at protocols.io.

## Background

In recent years, advances in long read sequencing technologies have enabled genome assembly to an unprecedented quality and quantity. These advances underpin the goal of the Earth BioGenome Project (EBP), which is to create high quality reference genomes for all described eukaryotic species [1]. This ambitious project faces many challenges from collecting and identifying species at scale, to extracting sufficiently high quality and quantity of DNA and RNA from a wide range of taxa, to sequencing, assembling and annotating extraordinarily diverse genomes. It is this central DNA extraction challenge that we address here, alongside sharing all protocols that enable our work. The EBP goal will only be met through open and rapid sharing of key protocols and pipelines.

The Tree of Life (ToL) programme at the Wellcome Sanger Institute is a major contributor to EBP goals. Over the past five years, we have extracted DNA and RNA from 41 phyla representing 4883 species under projects such as Darwin Tree of Life [2] and Aquatic Symbiosis Genomics [3]. We have released dozens of protocols at the Sanger Tree of Life Workspace on protocols.io to assist others in their efforts to carry out the laboratory work necessary to generate high quality reference genomes. These protocols for tissue preparation, high molecular weight (HMW) DNA extraction, fragmentation and clean-up, and RNA extraction have been applied at scale with standardised quality control (QC) measurements at key stages. Here we share both the routine processes that we employ as a first pass for organisms from a variety of different taxonomic groups as well as the approaches we take when we encounter failures. We also share things that we have learned along the way regarding specific challenges presented by different taxonomic groups, sample types, and species. The work presented here provides a summary of our first five years of work, with a frozen data set [4] used to enable us to provide success rates and review progress. Work in all of these areas is also currently ongoing, with new species being processed daily and new protocols developed to improve output and efficiency.

### Developing a standardised pipeline for the processing of diverse biological samples for reference genome assembly

The path from specimen to genome assembly requires optimal execution of complex wet lab, sequencing and informatic processes. In ToL, we use long read genomic sequencing and Hi-C chromatin conformation sequencing for reference genome assembly, and produce transcriptomic data through short read RNA-seq for primary annotation of completed genomes. Focussing on the wet-laboratory work, we have standardised the processes to allow progress at pace. In general, the laboratory steps required for generation of high-quality reference genomes are:

1. Sample Preparation: This step involves assessment of extremely diverse samples varying in size, density, morphology, and chemistry. Tissue typically will undergo some kind of homogenisation and be aliquoted for different pipelines (RNA extraction, DNA extraction, and Hi-C).
2. HMW DNA extraction: The protocols associated with this step comprise the most diverse set of protocols depending on the target taxon and the nature of the available tissue.
3. HMW DNA Fragmentation: Our primary long read sequencing approach over the past five years has been PacBio HiFi, and this requires molecules in the 12-22 kb range, which is shorter than typical HMW DNA extractions.
4. Fragmented DNA clean up: After shearing, it is important to perform a clean up to remove low molecular weight (LMW) DNA as well as compounds and inhibitors that may have co-extracted with or bound to DNA to achieve good sequencing results.
5. Hi-C: Samples are crosslinked to preserve the 3D structure of the genome, digested with restriction enzymes, biotin-labeled, and proximity-ligated before short read sequencing.
6. RNA extraction: RNA of sufficient quantity and quality for genome annotation is extracted and sequenced.

We have adapted and further developed protocols for the steps above from a wide range of primary sources. These protocols have been written in a modular way so they can each be used in conjunction with one another, depending on the taxonomy, tissue type and mass of the sample. They have all been published on protocols.io [5] in the Sanger Tree of Life Workspace [6], where we will continue to publish new protocols as we develop and deploy them. We encourage people who modify or improve these protocols to “fork” them on protocols.io and make them publicly available to the wider biodiversity genomics community. The datasets that have been produced during this work are provided as supplementary material [4], with one file containing data pertaining to the DNA extraction results, one to the DNA fragmentation results, and another one to the RNA extraction results as well as a data dictionary to facilitate interpretation. These files provide a more detailed view into the performance of various protocols on a wide range of species (e.g. there are nearly 5000 species in the DNA extraction results). As work continues, access to the ever growing data set has been made available via a searchable online ‘Portal’ at links.tol.sanger.ac.uk/datasets/tol-lab-data. All statistical analyses presented were performed using the statistical programming language R (Version 4.4.1 [7]), with data visualisations supported by Tableau [8] software (Version 2024.2.1).

The typical sample follows a four step path for HMW DNA extractions and processing (Figure 1), with samples branching off for Hi-C and RNAseq. First, a sample is examined and weighed and the taxonomy is noted using the information provided by the collector. Based on these features it is then directed into one of the three homogenisation protocols. The outcome of each of these protocols is three samples per species; two ‘tissue prep’ samples that can be directed toward any HMW DNA or RNA extraction protocol, and another sample to enter the Hi-C protocol. The processing of tissue to enter our Hi-C protocol differs depending on taxonomy (described in Table 1). Separately, HMW DNA is extracted from the prepared sample. Currently, we have ten HMW DNA extraction protocols, and one pre-extraction treatment each one optimised for different taxonomy and tissue types, detailed further below. All of our protocols are version controlled with the version number in the document name. Retired versions remain available on protocols.io but we advise using the most recent version number for any given protocol.

**Figure 1.**
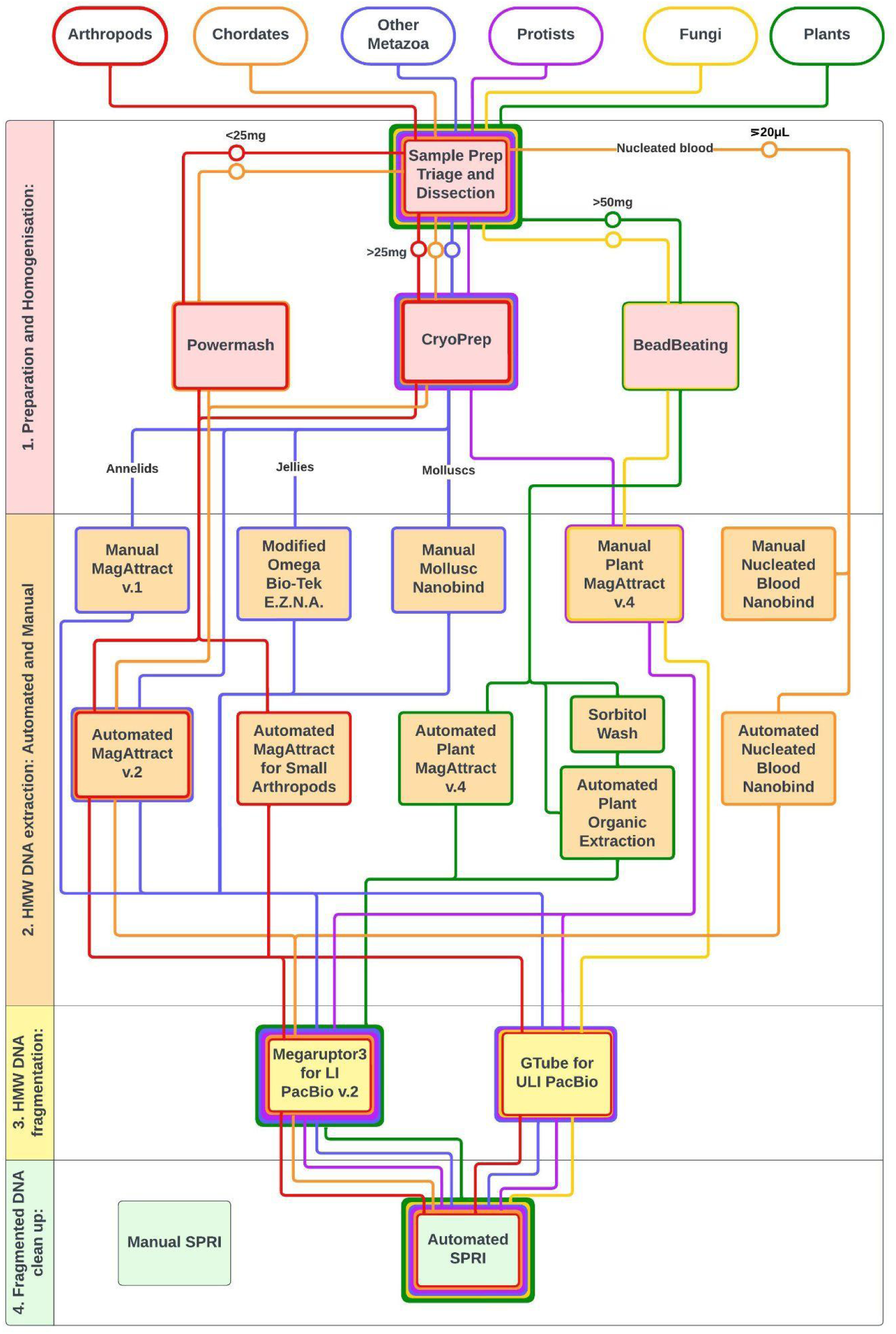
A tube map of Tree of Life protocols for HMW DNA extraction and processing. The publicly available ToL protocols for HMW DNA extraction and processing are categorised into the four numbered steps typically required to generate long read data for a high quality genome assembly. Each box here refers to a protocol available in the protocols.io Sanger Tree of Life Workspace and also linked to the Earth BioGenome Project Workspace. The current best practice is indicated by the route taken by samples from different taxonomic groups, shown by the coloured ‘tube lines’, and decision points to mark entry into these lines are discussed in the relevant taxonomic sections of this manuscript. The group ‘Other Metazoa’ includes mostly marine non-Chordata and macroalgae, and within ‘jellies’ are jellyfish and ctenophores.

**Figure 2.**
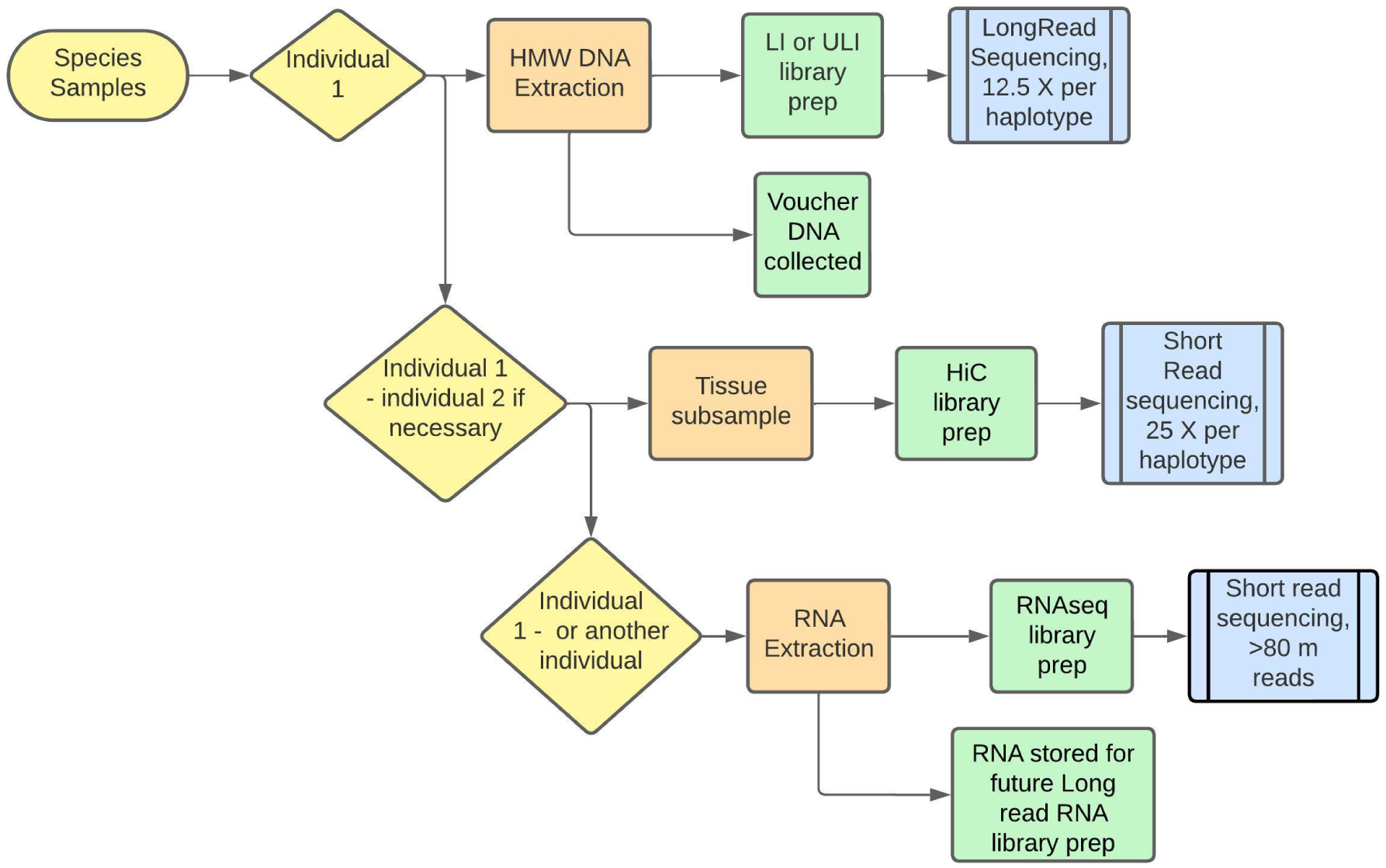
Process flow from species samples to the full data set required for genome assembly depending on tissue availability. Ideally, one individual specimen should provide tissue for generation of all data types. For many smaller organisms this is not possible, and a second or third individual may be required (indicated as individuals 1, 2 and another). Importantly, all long-read data for the initial contig assembly must be produced from one individual. If insufficient coverage is achieved from initial sequencing, this needs to be topped up with additional data generated from DNA from the same specimen. If there is very little DNA remaining, it may be possible to make a ULI library from what remains. Otherwise, long read data generation must start afresh from a new individual. The stated coverage follows the recommendations of the Tree of Life assembly pipeline at the time of writing.

**Table 1.** Sample Preparation Guidelines for RNA Extraction and Hi-C for each taxonomic group.

| Sample taxonomy | Ideal input for RNA extraction | Ideal input for Hi-C |
| --- | --- | --- |
| Arthropods | <10 mg powermash<br>>10 mg cryoPREP | whole head or up to 20 mg tissue,<br>no homogenisation |
| Chordates | <10 mg powermash<br>>10 mg cryoPREP | up to 20 mg tissue,<br>no homogenisation |
| Plants | ≥10 mg bead beaten | 50 mg bead beaten |
| Fungi | ≥10 mg bead beaten | 50 mg bead beaten |
| Protists | ≥10 mg bead beaten | 20 mg bead beaten |
| Other metazoa<br>and macroalgae | <10 mg powermash<br>>10 mg cryoPREP | up to 20 mg tissue,<br>no homogenisation |

The output of each of these HMW DNA extractions protocols is a sample that can be fragmented using either of two methods, the selection of which is dependent on the quality and quantity of DNA in the sample, and the intended PacBio library type. There are two main types of long read library preparation: the conventional Low Input HiFi library (LI PacBio) and the amplification-based Ultra Low Input HiFi library (ULI PacBio). The LI PacBio approach requires at least 500 ng of 12-22 kb fragment size DNA per Gb of genome (i.e. a 2 Gb genome would require at least 1 µg of sheared DNA). The ULI PacBio approach requires a shorter fragment size of around 10 kb to enable successful amplification, but can be successful with as little as 20 ng of sheared DNA for smaller (< 1 Gb) genomes.

Given these different input quantity and molecule length requirements, once we know the yield of DNA we have achieved in the HMW DNA extraction and its initial profile prior to shearing, together with an understanding of the predicted or known genome size obtained *via* GoaT (Genomes on a Tree database; https://goat.genomehubs.org/) [9], we choose the appropriate shearing approach. We use g-TUBES (Covaris, Woburn, MA) for ULI libraries and the Megaruptor (Diagenode, S.A.) for LI libraries. The output of these fragmentation methods can be submitted to either of the two clean up protocols depending on the scale of the operation (manual [10] or automated [11]), both of which are Solid Phase Reversible Immobilisation (SPRI) [12] methods.

Finally, RNA extraction is carried out on a separate tissue aliquot from the same species and where possible, the same organism. We deploy either a manual TRIzol protocol [13] or an automated MagMAX mirVana protocol (Thermo Fisher Scientific, UK) [14].

The modularity of these protocols allows for flexibility and a high throughput, while maintaining a standardised workflow. Having processed thousands of samples through these protocols, we have been able to monitor successes and failures that are both obvious (e.g. they yielded insufficient quality or quantity of DNA) as well as less obvious (e.g. where the DNA passed QC but still failed to generate sequence data). In practice, monitoring outcomes across diverse taxa has enabled us to generate reference level genomes at pace for those taxonomic groups that tend to yield good DNA and results with the protocols above, while also highlighting taxonomic groups that do not proceed well and thus require further attention and R&D protocol development.

### Sample selection and data generation in an ideal world

In an ideal process, all data for a species’ genome assembly would be generated from the same individual, avoiding issues of sequence diversity among individuals. However, this is often not possible due to limited tissue availability. Assembly algorithms work best when all long-read data for the contig assembly is produced from a single individual, therefore we always aim to access sufficient DNA to preclude needing to start long read processes over with a new individual. Generation of Hi-C and RNAseq data from the same individual as was used for long read data generation is optimal, but for these processes, using different individuals is viable. When two or more different individuals must be used for data generation, ideally data from long read and from Hi-C should be from the same sex such that the Hi-C data represents the full complement of chromosomes present in the long read data. Any individual can be used to produce transcriptomic data, and in instances where several individuals are available it is possible to start all of these lab processes in parallel. Individuals should be selected bearing in mind their biology, e.g. the heterogametic sex and non-polyploid samples are preferred following EBP guidance [15]. When specimens are large enough for dissection, or where multiple tissue types are available for a species, different tissues can be selected for different processes. For example, in insects, we would usually generate long read data and Hi-C from the head and thorax, and only use the abdomen for RNAseq if necessary. This avoids sequencing the microbiome present in the gut, and, in parous females, any sperm or embryos present in the reproductive tract. Our general rule is to avoid tissues that might contain organisms in addition to the target species. While it is interesting to assemble the cobionts in a sample, the additional sequencing data required, and the complexity of the subsequent assembly task argues against these tissues as sources for genomic DNA isolation.

The amount of long read data required for genome assembly is dependent on genome size and is typically described in terms of coverage (e.g. 25x coverage of a diploid 1 Gb genome = 25 Gb of data required to give 12.5x coverage per haplotype). These calculations increase where polyploidy is present, as 12.5x coverage per haplotype is the minimum required. For this reason, the predicted haploid genome size and ploidy for a species is retrieved from GoaT [9] to help determine the initial amount of sequencing required. For most species a directly measured genome size is not available and estimates from an average of the nearest taxonomic neighbours are used. Whilst these estimates can be inaccurate, they provide a reasonable starting point for sequencing efforts, which can be adjusted based on k-mer based genome size and ploidy estimates obtained from initial data.

### Sample Preparation

Most samples are provided as small pieces of tissue, cell culture pellets, or whole small organisms, snap frozen at collection in 1.9 mL FluidX (barcoded) tubes, transported and stored at −70°C, in line with EBP guidance [15]. Where tissue is abundant, such as vascular plants, larger tubes (7.6 mL) are used to collect as much tissue as possible without compromising the integrity of the tissue. Once a sample has been selected for work in the laboratory, a process is followed with the aim of normalising the biologically diverse samples as much as possible, resulting in the production of a tube containing sufficient material for the next downstream process. The ideal amount of starting material is usually 25 mg for animals, protists and fungi and 50 mg for plants. Despite the fact that many organisms weigh less than 25 mg in total, we progress these through the protocols and are often successful.

All samples are weighed and divided based on their taxonomy, tissue type and size/mass of material for disruption following the sample triage protocol [16]. Tissue homogenisation is a crucial step prior to HMW DNA extraction. We have used the Powermasher (Nippi, Japan), cryoPREP (Covaris, Woburn, MA) and FastPrep-96 (MPBio, CA) at scale following the guidelines set out in Table 1. In general, smaller samples are weighed and then powermashed [17] in the extraction lysis buffer at room temperature. The benefits of this method are the ability to adapt the duration of the treatment to the requirements of the sample structure, and directing the disruption toward different parts of the sample as it disrupts, i.e. concentrating on more resistant pieces of tissue as they become apparent during the process. Importantly, there is no loss of tissue since the process occurs within the lysis buffer, and all material is immediately put into nucleic acid extraction without any tube transfer or pipetting. The drawback of this technique is its low throughput nature, with each sample requiring individual powermashing.

Homogenisation at extremely low temperatures can be achieved using a pestle and mortar, and liquid nitrogen. This approach is inherently low throughput and it can be hard to avoid cross-contamination of samples through residual tissue on instruments. The cryoPREP instrument [18] (Covaris, Woburn, MA) solves the issue of cross contamination. Samples are placed into proprietary bags (TissueTUBEs) made of material resistant to extremely low temperatures and force (Figure 3). The whole bag containing only the tissue sample (no lysis buffer) is submerged in liquid nitrogen and then placed on the machine to be smashed between metal plates. The cryoPREP can be used repeatedly on the same sample, and the strength is adjustable. Therefore, the process can be continued until the sample is reduced to a fine powder. The bag can be repeatedly submerged in liquid nitrogen between pulverisations, maintaining low temperature and preventing degradation of nucleic acids within the sample due to the action of endogenous nucleases. We note that repeated processing can become labour intensive on the cryoPrep, and when processing a large number of samples maintaining the cold temperature is a challenge. Additionally, the proprietary bags cannot be reused and add significant cost to sample prep.

**Figure 3.**
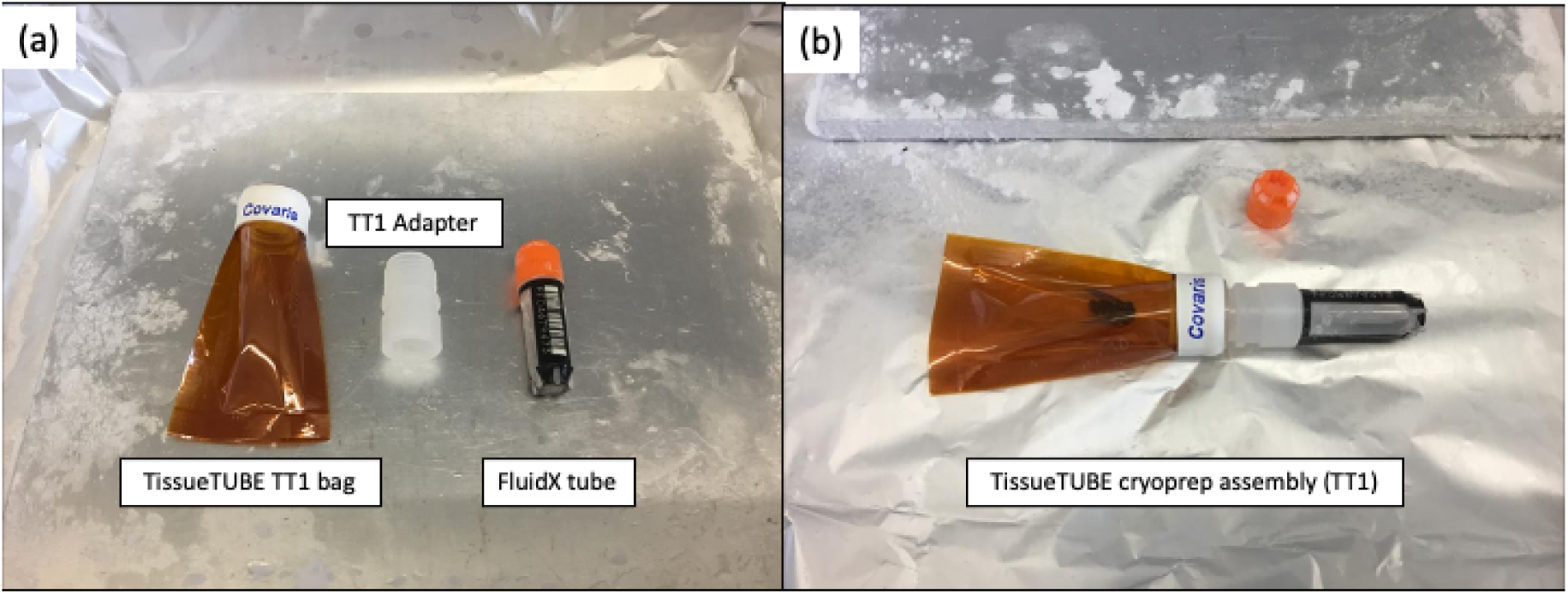
TissueTUBE assembly for the cryoPREP. (a) Left to right: Covaris TT01 TissueTUBE, TT01/1.9 mL FluidX adapter and 1.9 mL FluidX tube. (b) assembly with sample in place. The sample is stored within the FluidX tube and once all parts are assembled, the assembly is inverted to allow the sample to move into the TT1 bag. The TT1 Adapter shown was 3D printed within the Sanger Institute.

The FastPREP-96 bead beating approach is useful for plant tissue disruption [19]. FluidX tubes containing snap-frozen dry tissue samples (50-90 mg) are selected, and 3 x 3 mm stainless steel grinding balls added. Up to 48 of these sample tubes are then assembled into a rack which is submerged in liquid nitrogen to cool, and then carefully lifted allowing any excess to drain. This chilled rack of tubes is then placed on the FastPrep-96 instrument to be shaken at 1600 rpm for 30 seconds. This process, including the submersion, is repeated three times, after which all samples are reduced to a homogenous powder. There is no need to remove the beads from the tube before starting the lysis process, and performing this in the same tube prevents loss of tissue. Two racks of tubes can be processed in parallel, enabling a throughput of 96 samples. This technique has proven extremely successful for plant tissue disruption and is showing promise for other organisms and tissue types. For a set of test species (oak, ladybird, snail, yeast, and marine fungus), tissue disruption with FastPrep-96 achieved similar results as cryoPREP and powermasher, but with the processing advantage of scale and a standardised approach.

While figure 1 shows the ideal sample weight and homogenisation method for our HMW DNA protocols, Table 1 shows the ideal preparation for RNA and Hi-C, each split by taxonomic grouping. The ideal preparation of samples is dependent on the process for which the sample is intended, which can make standardising decisions difficult when balanced against the diverse nature of the samples. For some sample types, such as protists, the ideal method of disruption has not yet been ascertained so the current best practice is shared here. Details are provided in the taxon specific sections of this manuscript to further describe the observations that can be drawn from our work so far. For other groups such as chordates, arthropods and plants, our methods are robust and routine but the FastPrep-96 approach may replace powermashing and the cryoPREP and testing is underway.

### High Molecular Weight DNA Extraction

Following the sample preparation and appropriate disruption process, samples progress to HMW DNA extraction. The ideal input weight and disruption method for different sample types is shown in Figure 1. A proportion of samples do not meet the minimum mass criteria, and it is therefore not possible to standardise the input for these samples. This does not prevent these samples from entering the process and contributes to our understanding of performance outside the ideal parameters.

In order to minimise the number of different extraction protocols we use, our approach has been to first test a sample from every species using one standardised protocol. Samples that pass well through this extraction protocol will go forward to produce sequence data, and those that fail highlight the species groups that require further investigation. We use the Qiagen (Hilden, Germany) MagAttract HMW DNA extraction method [20] as our default first protocol due to the track record seen in the laboratory and the ability to automate on the KingFisher Apex [21]. For plant samples, an accompanying Plant MagAttract protocol [22] has been implemented. Because the samples received are diverse, if a first extraction fails a second attempt is made with the same protocol. This allows for the selection of a new individual, or a different tissue type, and on many occasions this results in a successful DNA extraction. This may be due to many factors including individual differences within species, or to factors relating to the sample collection and preservation. A 10 µL molecular voucher (aliquot) of every DNA extraction performed is retained for deposition to museums, and in some cases, this voucher can be used in part for “top up” when slightly more long-read coverage is required.

Standard quality metrics are collected for each sample after DNA extraction. Nucleic acid quantity is measured using the Qubit® dsDNA assay (Thermo Fisher Scientific, UK). We also assess DNA purity through spectrophotometry using a Lunatic spectrophotometer (Unchained Labs, Pleasanton, CA.). We measure the ratio of absorbances at 260 nm:280 nm, which is ideally ∼1.8, and 260 nm:230 nm, which is ideally between 2.0 and 2.2. Deviation from the optimum for either of these measures indicates the presence of contaminants in the extraction, for example phenols or carbohydrates, that may interfere with downstream processes. The fragment length distribution of the HMW DNA is assessed using the FemtoPulse System (Agilent Technologies, Santa Clara, CA.) and their Genomic DNA 165 kb Kit. This pulsed-field capillary electrophoresis system measures concentration (through spectrophotometry) and length (based on retention time relative to standards) of the extracted DNA, and provides accurate sizing of fragments up to 165 kb. Above this size, ultra HMW fragments are visible, but the sizing is not accurate.

The quality of DNA extracted from samples is highly variable, resulting in diverse FemtoPulse profiles. To assess the traces routinely and standardise decision making between users, we developed a categorisation system. We defined five profile classes: “LMW DNA”, “smear bulk <50 kb”, “smear bulk >50 kb”, “HMW band plus smear” and : HMW band”. Model profiles representing each of these categories is shown in Figure 4. Over time, as we become more familiar with certain taxonomic groups (such as Lepidoptera), some samples are processed as scale without the routine labelling of profiles [4]. For more challenging sample groups still under active R&D, different aspects can be noted using a multi-select approach - for example “HMW band” and “LMW DNA” could both be selected for one sample [4].

**Figure 4.**
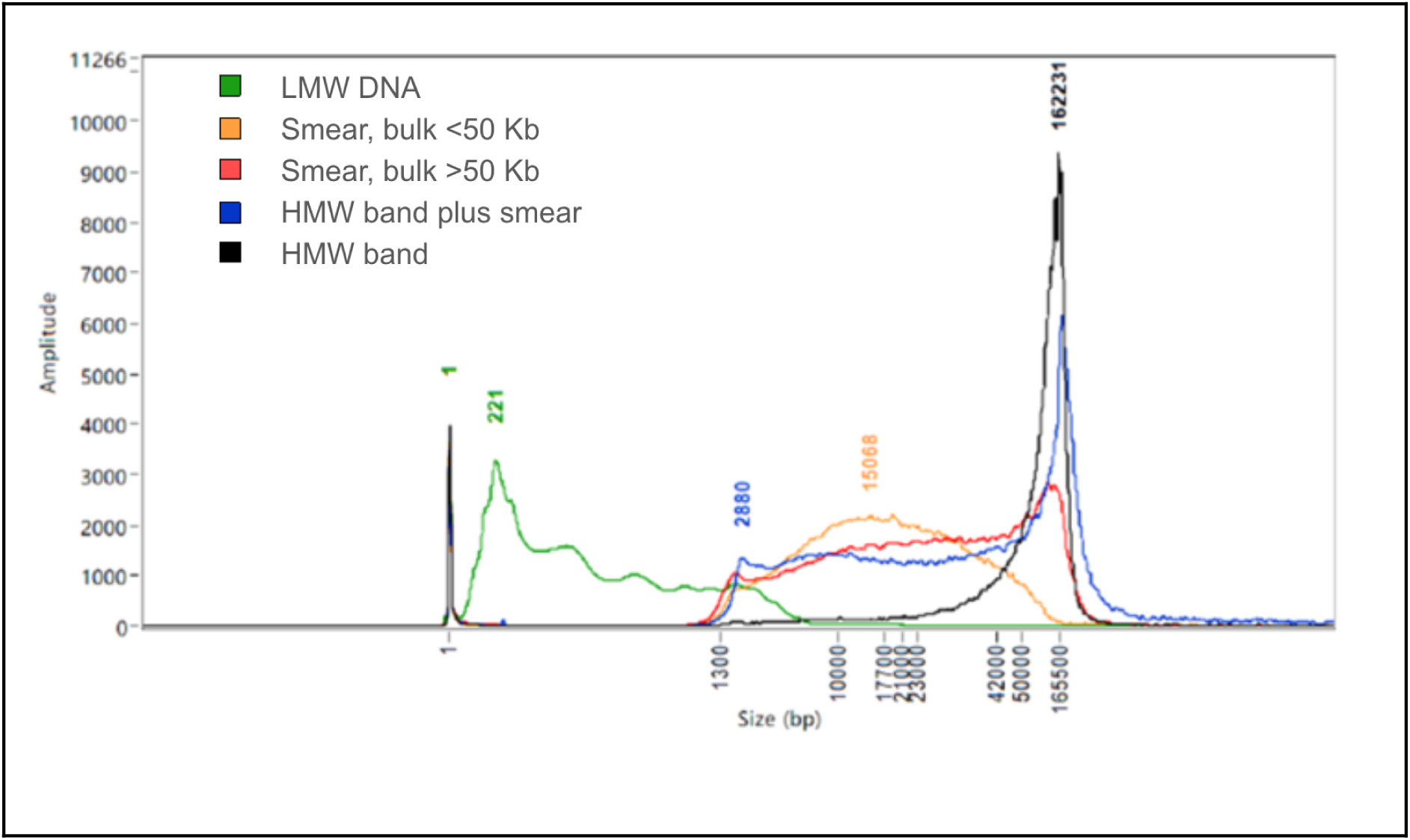
Representative FemtoPulse molecular weight profiles of five DNA extractions, modelling the categories used within the Tree of Life. The X axis shows fragment size. Ideally most DNA would sit above 50 kb. This is demonstrated in the example traces for categories ‘HMW band’ (black), ‘HMW band plus smear’ (blue) and ‘Smear bulk >50 kb’ (red), where peaks in the trace are visible at approximately 160 kb. The profile ‘Smear, bulk <50 kb’ (yellow) is common and can be progressed best when it is possible to remove the smaller fragments - ideally removing everything below 10 kb. Finally, the category ‘LMW DNA’ (green) is a failure for downstream long read sequencing.

In case of good DNA quality but low yield on first extraction, there are two options. In some cases, it may be possible to extract again from the same specimen and pool samples in order to achieve the required quality and quantity of DNA. If this is not possible, samples with < 1 Gb predicted genome size yielding over 100 ng of HMW DNA sufficient quality are routed toward ULI library prep, aiming for >20 ng of DNA after shearing and SPRI cleanup. Samples with <100 ng of DNA may be progressed along this route if there is no option to repeat the DNA extraction, i.e. there is no tissue remaining. With the new PacBio Ampli-Fi kit, these thresholds are likely to be 1 ng per 3 Gb genome size. For species where even this quantity of DNA is not achievable, picogram-input methods like PiMmS are available [23]. The ULI option is restricted to species with a genome size of <1 Gb due to the impact of amplification bias and the resulting poor coverage of specific genomic regions that affects downstream assembly quality.

We have introduced a 0.45X SPRI step directly after extraction to remove DNA fragments <10 kb [24]. Because the FemtoPulse trace shows relative absorbance normalised to the maximal value, it can be difficult to assess the fragment distribution for samples with significant amounts of LMW DNA. The removal of shorter fragments via the SPRI cleanup enables more accurate analysis of the HMW DNA profile, demonstrated by analysis of *Martes martes* (pine marten) samples (Figure 5). Improving the analysis and decision making process at this point enabled a reduction in the quantity threshold for passing extractions through to fragmentation.

**Figure 5.**
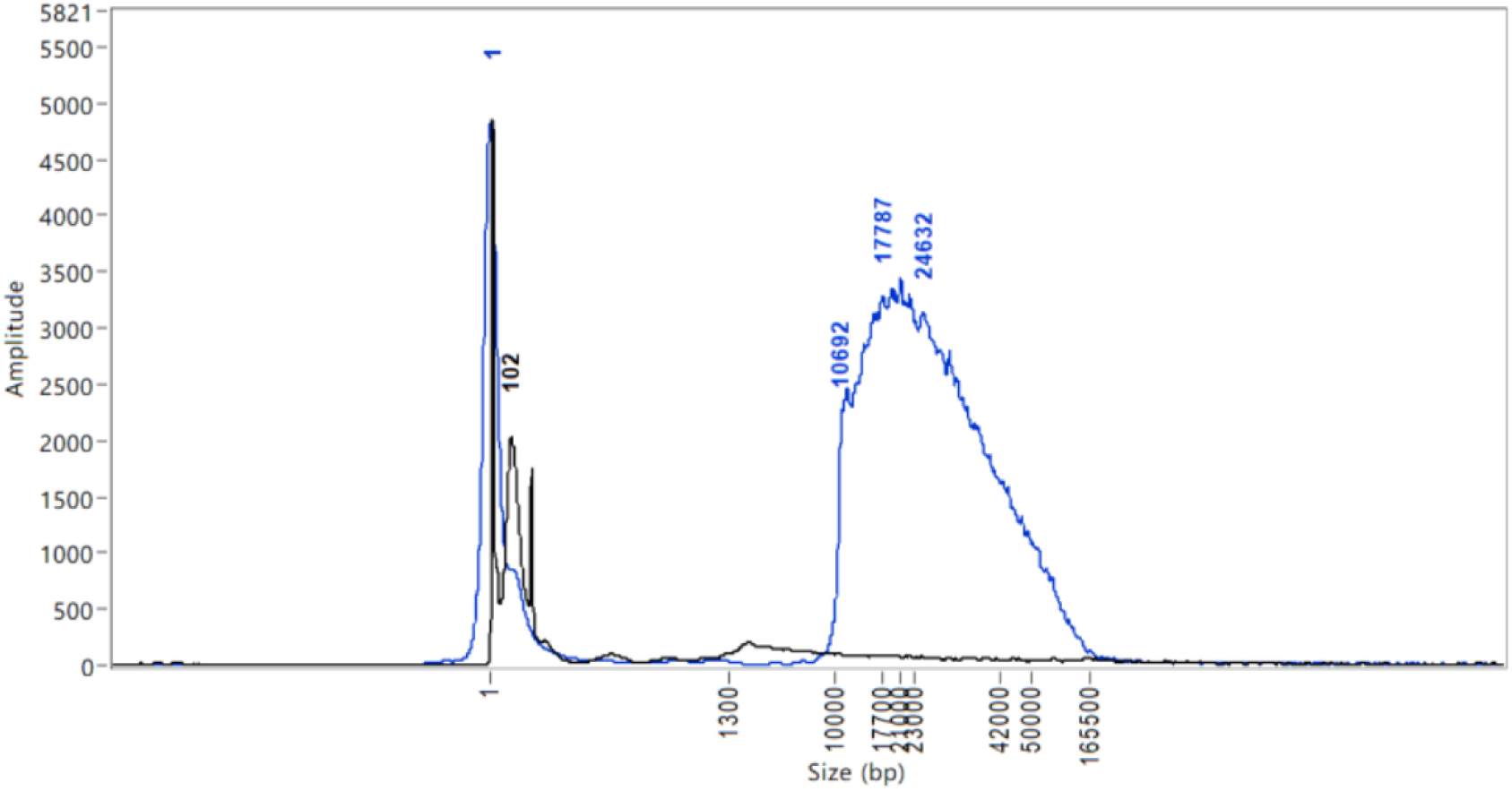
Overlaid FemtoPulse profile for DNA extractions from *Martes martes* heart tissue (Mammalia; pine marten). The overlaid profiles show the impact of performing a 0.45 SPRI after DNA extraction. The black trace (manual DNA extraction with no SPRI), shows very little detail due to the large LMW peak, whilst the blue trace (automated DNA extraction with a SPRI) reflects the profile of the remaining DNA with significantly more detail allowing for informed decision making.

The overall success rates of samples within laboratory extraction processes can be broken down by taxonomic group (Figure 6). Overall we find that chordates and plants progress well, showing the highest HMW DNA extraction pass rate of 96% (91.2 Pass, 4.0% Pass ULI, 1.1% Pooling), and 91% (84.3% Pass, 5.5% Pass ULI, 0.9% Pooling), respectively. The number of species within the arthropods dwarfs the other sample groups, with 2373 species having been processed with a total pass rate of 85%. The highest HMW DNA extraction fail rate is observed in fungi, at 34.2%.

**Figure 6.**
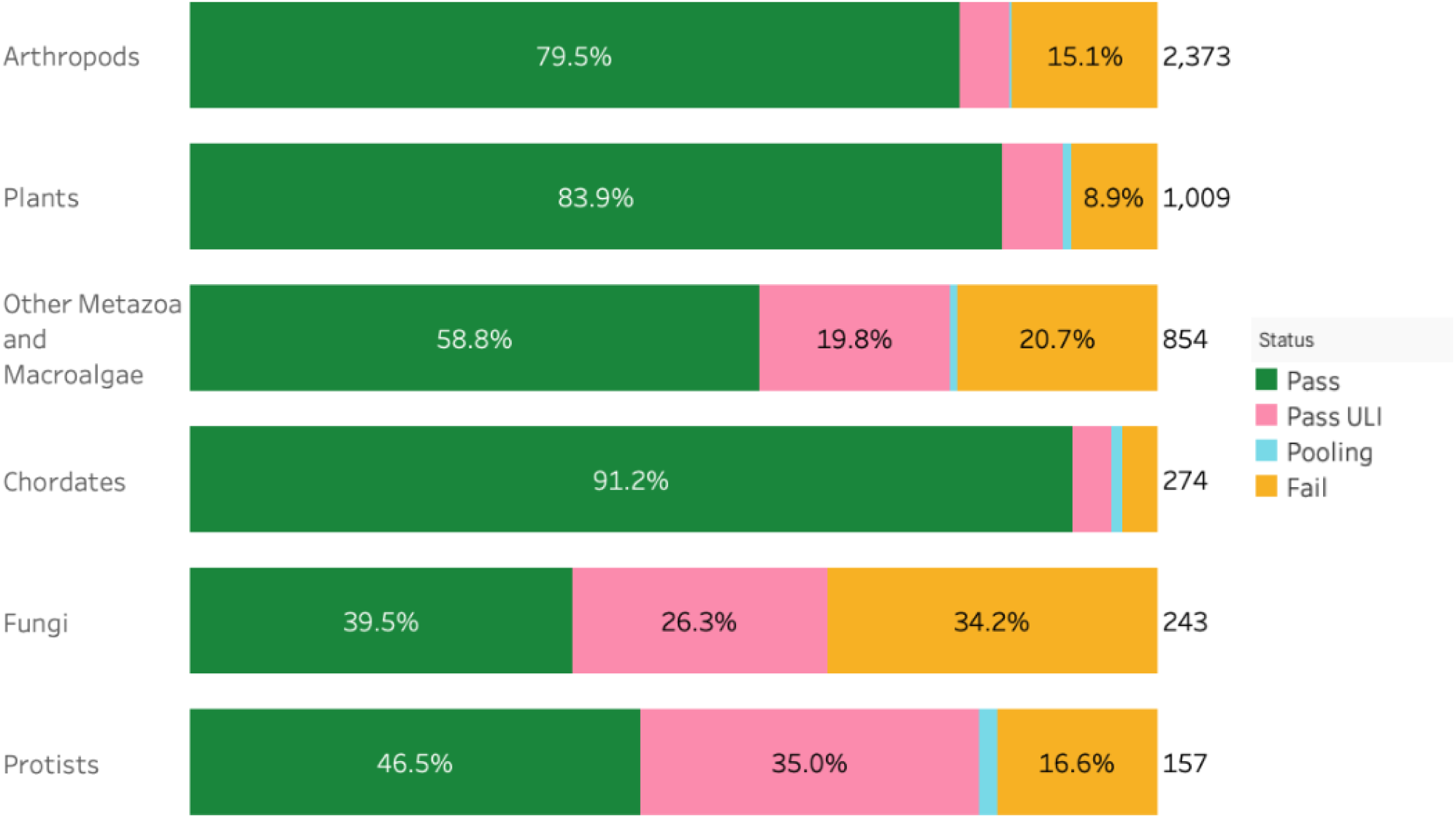
Success rates of DNA extractions per species from six taxonomic groups. The bar chart summarises the DNA extraction success per species across the six taxonomic groups. The results are categorised as: Pass – DNA sufficient for sequencing achieved; Pass ULI - DNA sufficient for sequencing with ultra low input achieved; Pooling - two DNA extractions were pooled to meet QC threshold; Fail - extractions have failed to provide sufficient quality and/or quantity DNA to proceed. The results represent the best DNA extraction outcome per species, determined using the hierarchy: Pass > Pass ULI > Pooling > Fail. The number to the right of each bar represents the total number of species processed within each taxonomic group, and the percentages inside the bars indicate the proportion of species in each category.

### HMW DNA fragmentation

#### Megaruptor fragmentation

The DNA fragmentation protocol [25] using the Megaruptor 3 (Diagenode, S.A.) instrument, forces extracted DNA through a single-use hydropore connected to a syringe at a controlled rate, enacting mechanical shearing upon the DNA within the solution. The system can be used at various speeds, producing fragments of different median length. For LI PacBio libraries the sheared DNA lengths should be in the range of 12 - 22 kb, a tight peak (e.g. most DNA at 18 kb). The main challenge with Megaruptor shearing is pore blocking, where a syringe fails to pull the sample back through the hydropore after the initial pass. This occurs without warning and is largely unpredictable, although it is more common with visibly viscous DNA extractions. One way to overcome blockage is by diluting samples and running portions of the sample through the syringe over multiple stages, or transferring the extract to a different hydropore syringe designed for viscous samples (Cat. No. E07020001). Occasionally a sample does not shear fully during this process, which is clear only after observing the fragment size distribution on the FemtoPulse. A second attempt at shearing is then made which typically completes the shearing, however, if a minority of longer fragments persist the samples are still progressed to library prep where they will be removed in later size selection processes.

#### g-TUBE fragmentation

The incorporation of DNA amplification into the ULI library prep method allows for a significantly lower input (as little as 20 ng post shearing), and a shorter fragment length of 9-11 kb. This fragment length can be routinely and reliably achieved using a g-TUBE (Covaris, Woburn, MA) with our standard protocol [26], which relies on shearing due to forcing the DNA through a narrow aperture membrane. While Megaruptor shearing can also be used to generate the smaller fragments needed for ULI, g-TUBEs are preferable because of faster processing time and more reliable output. Unlike the Megaruptor syringes, the g-TUBE only requires the use of a microcentrifuge for shearing. Occasionally a sample fails to pass through the g-TUBE and requires a second spin, but this is the extent of troubleshooting required for this method.

### Fragmented DNA clean up

After fragmentation with either method, the DNA is cleaned and concentrated again using SPRI beads, either manually [10] or automated on the Kingfisher APEX [11]. This process both purifies the DNA and removes shorter fragments. Following this, the DNA is evaluated using Qubit and Lunatic spectrophotometry and FemtoPulse electrophoresis. Samples meeting the criteria for LI or ULI sizes and yields progress through to PacBio library preparation.

The QC results inform a decision-making process, resulting in an output of either a ‘Pass’ or ‘Fail’ post-shearing. The pass rates for shearing are relatively high – from 89% for chordates to 67% for arthropods (calculated from data in [4]). The ULI path provides a route for low yield samples, and also those that fail because their short fragment length is not ideal for LI. Protist samples exemplify this, with a pass rate of 20% in LI fragmentation and 54% in the ULI method (calculated from data in [4]). DNA samples that fail in both shearing options would require another DNA extraction event. Since the introduction of a SPRI after DNA extraction and before fragmentation, there has been an increase in the fragmentation pass rate. Samples that would previously have failed at this point are now removed earlier in the process, meaning that less time is spent processing samples that are not suitable for sequencing.

### Long Read Sequencing

Ultimately, the success or failure of a HMW DNA extraction for the purposes of long read sequencing is judged by sequencing yield, read quality and read length. For the PacBio platforms, raw sequencing reads from circular library molecules that contain multiple reads across both strands of the insert DNA are automatically error corrected to generate circular consensus sequencing (CCS) output reads of high per-base quality. On the Sequel IIe platform, a CCS yield over 20 Gb is considered good, while a yield of 10-20 Gb is not ideal but still potentially adequate depending on genome size. Finally, a yield below 10 Gb is considered poor and a target for improvement. The Revio platform is designed to yield three times this output, and performance is judged in line with this. With the aim of producing 12.5x coverage per haplotype, library multiplexing on the Revio is advised unless work is taking place on large genome organisms (e.g. > 2 Gb) to avoid overproduction of data.

If a sample is sequencing well but has not reached the required coverage, a ‘top-up’ of data from the same genetic individual is needed. Recent work in this area has shown that the longevity of both LI and ULI libraries is greater than had been anticipated, with examples of both surviving storage at −70°C for over 9 months before performing equally well on a second run. Where no library or DNA remains, the DNA voucher (a 10 µL aliquot taken from all extractions) can be valuable for ULI prep.

Small diploid genomes (<0.5 Gb) can reach 25x coverage from low cell yield (i.e. <10 Gb), but this poor performance remains a target for improvement. As the data are collected and accumulated, trends in lower CCS yield for different taxonomic groups become indicators of R&D need.

The data yield required for successful assembly is based on the genome size of the target organism within a sample. However, as samples are collected from wild environments other species are often present within a sample (e.g. the microbiome, pathogens and parasites). The bioinformatics pipelines in place to process data are capable of filtering out data that derives from these “cobionts”. In many cases the reads from non-target organisms are sufficient to generate cobiont genome assemblies as a by-product of the attempts to sequence the target species. However, occasionally non-target species can be present in such abundance as to prevent the sequencing of the target species or make it too costly to continue sequencing to achieve required coverage for the target species. This scenario may arise from cultured species that require the presence of other organisms to grow, for example protists that feed on bacterial species. These cases present a challenge for reference genome pipelines and purification of the sample upstream of DNA extraction is recommended.

The output from the standardised workflow described results in a range of CCS yields that shows variation for each taxonomic group for both ULI and LI submissions (Figure 7). We compared data production for all libraries run as one species’ library per cell on both PacBio platforms, Sequel IIe and Revio. To present standardised results, data from multiplexed samples were not included. Comparing submission types, the average yield of runs on the Revio instrument for LI libraries ranged between 48 and 69 Gb, and from 65 to 74 Gb for ULI submissions. On the Sequel instruments, the yield ranged between 17 and 25 Gb for LI submissions, and 19 to 24 Gb for ULI submissions. Arthropod species had a fairly consistent yield regardless of instrument or library preparation techniques. However, fungi had more variable yields, with low average yields of 17 Gb with the LI library prep method but much improved average yield of 23 Gb when ULI libraries were sequenced, on the Sequel IIe platform. Our experience with sequencing fungi on the Revio is limited but in line with the Sequel IIe, with approximately three-fold higher yields for LI libraries (50 Gb). Most fungal species are now directed to the ULI library pipeline because of low DNA yields. Overall, ULI libraries show less variation in yield within each taxonomic group than LI libraries, as would be expected for PCR-amplified DNA when compared with native.

**Figure 7.**
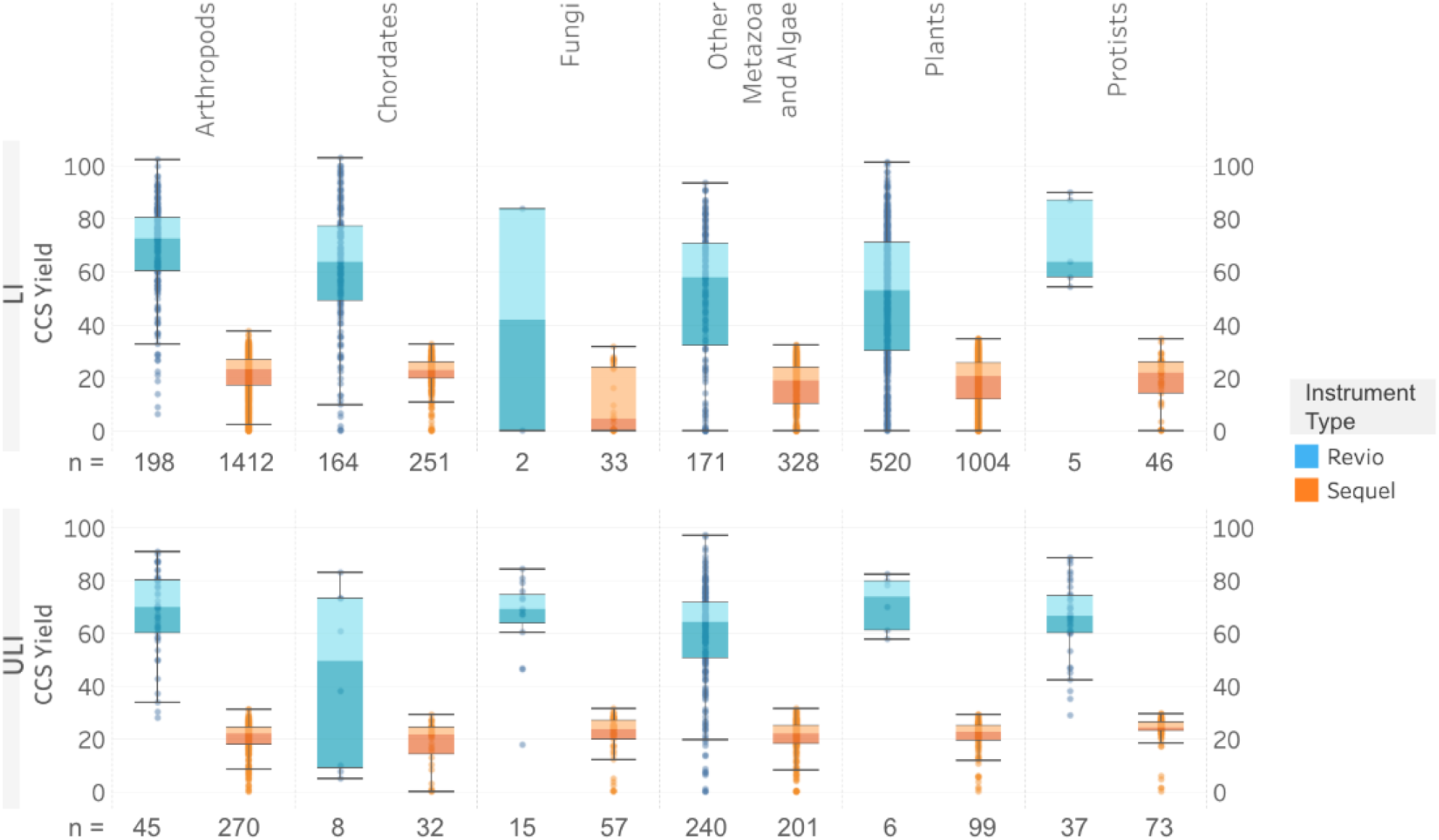
Distribution of CCS yield per taxonomic group. The distribution of the CCS yields across taxonomic groups is shown for each instrument type (Revio and Sequel) and library type (LI: Low Input; ULI: Ultra Low Input). Each box spans the interquartile range, with the lower and upper edges indicating the 25th and 75th percentiles, respectively. The horizontal line within each box represents the median CCS yield value per taxonomic group. Whiskers extend to the furthest data points within 1.5 times the interquartile range, while data points outside of this range are considered outliers. Data analysed included all single specimen libraries (i.e. non-multiplexed) sequencing runs, regardless of extraction method. The label under each subplot refers to n, the number of sequencing runs within each sub-category.

In addition to the CCS yields varying across species, they can also vary within a species. For example, in the case of the newt, *Lissotriton vulgaris*, which has a large genome (24 Gb), a single library was made from DNA extracted from muscle. This library was run on seven Sequel IIe cells, with CCS yields ranging from 18 to 32 Gb per cell. This variation and unpredictability in yield presents challenges for scaling production. The recent introduction of SPRQ preloading normalisation on the Revio system will, we hope, reduce this unwanted variability. To more accurately target required coverage based on predicted genome size, many libraries are now multiplexed, with 2, 4 or 8 libraries run on one Revio cell in parallel. Ideal plexing in terms of molarity, taxonomy and fragment length is not possible as each specimen varies in its final library insert size profile.

### Hi-C Library prep and sequencing

A tissue aliquot to be used for Hi-C library prep is created for each species during the sample preparation process. The guidelines for the amount and disruption of each sample type is shown in Table 1. We use the Arima Genomics (Carlsbad, CA, US) Hi-C v2 kit. Three distinct fixation protocols are used depending on the taxon group. For animals, we follow the Arima high coverage kit recommendation for animal tissue, which involves fixation by 2% formaldehyde for 20 minutes in TC buffer (Arima). For plant and algal samples, we follow the Arima high coverage recommendation for mammalian cell lines, which involves nuclei isolation using the Qiagen Qproteome Cell Compartment kit followed by fixation with 2% formaldehyde for 10 minutes in 1x PBS buffer. Finally, for sponges (Porifera) that have been prepared via the “squeeze” method [27] to create a cell pellet, we carry out fixation with 2% formaldehyde for 10 minutes in 1x PBS buffer.

Hi-C is performed according to manufacturer’s recommendations except that the number of PCR cycles used in Illumina library amplification is directed by the DNA concentration post adapter ligation and streptavidin enrichment as measured using Qubit dsDNA high sensitivity kit (Thermo Fisher Scientific, UK), rather than determining amplification cycles by qPCR as in Arima QC2 procedure. The following PCR cycle guidelines are used: If >8 ng/µL DNA in post streptavidin enrichment quantification use 8 cycles of PCR; If >2 ng/µL DNA in post streptavidin enrichment quantification use 10 cycles of PCR; If >0.5 ng/µL DNA in post streptavidin enrichment quantification use 12 cycles of PCR; If >0.1 ng/µL DNA in post streptavidin enrichment quantification use 14 cycles of PCR; For lower concentrations use 16 cycles PCR.

Libraries are sequenced using Illumina (San Diego, CA, US) short read technology on the NovaSeqX, 150 B paired end reads on the 25B flow cell. Libraries are multiplexed such that 25x coverage per haplotype of the genome is aimed for for each sample. Grouping together samples with genome sizes of <1.5 Gb, 1.5-2.5 Gb, and >4 Gb can be useful for achieving desired plexing levels.

### RNA extraction

For all species we also extract and sequence mRNA to provide data for gene annotation. These data would ideally be produced from several different tissue types for each species, as this is most beneficial for gene annotation, but this is often not possible due to small organism size or restricted number of tissues collected. Originally, a manual TriZol method [13] was applied that achieved a high success rate from an extremely wide range of samples, typically using 25 mg of tissue. Tissue prepared by either the cryoPREP or powermasher can be used as input for this method, and the resulting yields were consistently significantly greater than requirements for short read RNAseq (Illumina, San Diego, CA.). After isolation, any DNA remaining in the samples was removed using Turbo DNase (Thermo Fisher Scientific, UK) and RNA was checked for quality and quantity using the Qubit RNA Broad Range Assay kit (Thermo Fisher Scientific, UK) and the Nanodrop. This extraction method is ideal for a small number of samples, but the ergonomic issues and use of hazardous substances are prohibitive for scaling up. We therefore switched to the MirVana (Thermo Fisher Scientific, UK) bead-based extraction protocol [14] and reduced the amount of tissue input from 25 to 15 mg. All taxon groups score near to 100% extraction success, with the exception of protists at 92% pass rate (calculated from data in [4]), meaning a total RNA yield over the 100 ng input requirement for our standard library prep and sequencing process; Poly(A) RNA-Seq libraries constructed using the NEB Ultra II RNA Library Prep kit, following the manufacturer’s instructions, sequenced on the Illumina NovaSeq X instrument. For samples that fail, a different individual, different tissue type(s) and/or increased input amounts can be used in order to increase the RNA yield or quality obtained. Ultimately, RNA is highly dependent on the quality of the sample material provided and many failed extractions originate from samples not preserved in the ideal way. RNA extracted from several different organisms and tissue types using this method has also been successful for long read RNA sequencing with the Kinnex (Pacific Biosciences, Menlo Park, CA.) methodology.

### Taxonomic specific considerations

#### Arthropods

Small arthropod species are often preserved as whole individuals, requiring several individuals to complete the data required for an assembly (one for long read, one for Hi-C, one for RNAseq). Larger arthropods are partitioned into different tubes, e.g. head, thorax, and abdomen each in separate tubes. Arthropods have an extraction pass rate of almost 85% across 2374 arthropod species reported on here representing 1575 genera, 453 families, and 52 orders (Figure 8).

**Figure 8.**
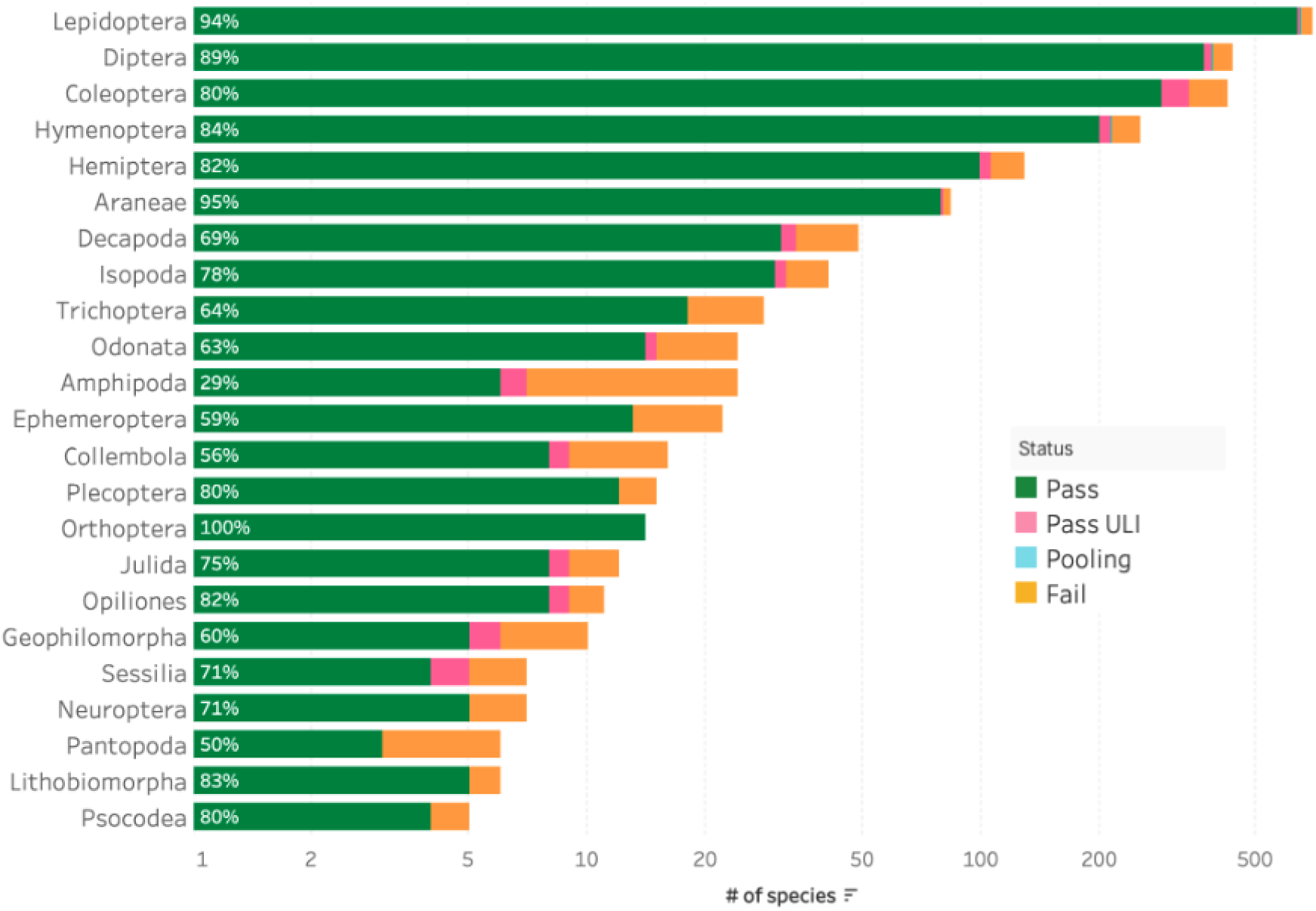
Arthropod DNA extraction success metrics by order. The bar chart summarises the DNA extraction success per species across Arthropod orders. The results are categorised as: Pass – DNA sufficient for sequencing achieved; Pass ULI - DNA sufficient for sequencing with PacBio Ultra Low Input achieved; Pooling - two or more DNA extractions were pooled to meet QC threshold; Fail - extractions have failed to provide DNA of sufficient quality or quantity to proceed. The results represent the best DNA extraction outcome per species, determined using the hierarchy: Pass > Pass ULI > Pooling > Fail. The number inside each bar represents the percentage of species that have passed extraction within the orders, including Pass, Pass ULI and Pooling categories. To account for the wide range in species counts, a logarithmic scale is used, and orders with fewer than five species are excluded from this visualisation but are available in the supplementary material (Figure S1).

Whilst arthropods usually perform well at extraction they are not without challenges. One challenge is the disruption of small organisms with chitinous exoskeletons, such as Amphipoda where 71% of 24 species have failed extraction (Figure 8). Initial work to apply the bead beating homogenisation method for these samples looks promising. Another challenge is small body size resulting in the most common failure being low HMW DNA yield. If the genome size is appropriate, these samples can be successful with ULI.

However, ULI does not work well on its own for small organisms with large genomes. Jumping spiders (Salticidae) are an example of such a group, with small sized bodies, typically 10-15 mg but ranging down to 3 mg, and large genomes of up to 10 Gb. A modified extraction protocol [28] with reduced volumes of buffers was successful in improving DNA yields per specimen.

Isopods are another example of species with typically small body size (<1 cm) and large genomes (e.g. *Oniscus asellus* 8.4 Gb). When tissue is restricted due to the organism size, and the DNA sequences poorly, as observed for isopods, reaching sufficient coverage is challenging. Currently, this challenge is being addressed through combining LI and ULI library types. This strategy minimises the impact of amplification biases present in the ULI data, as regions of drop out are likely to be compensated by presence in the LI data. The libraries produced from the ULI approach tend to sequence very well, as the DNA has been amplified. The abundance of DNA in relation to any inhibitors present is also changed in favour of high sequencing yields.

Arthropod DNA tends to perform well in fragmentation processes as shown by the high pass rates (Figure 9).

**Figure 9.**
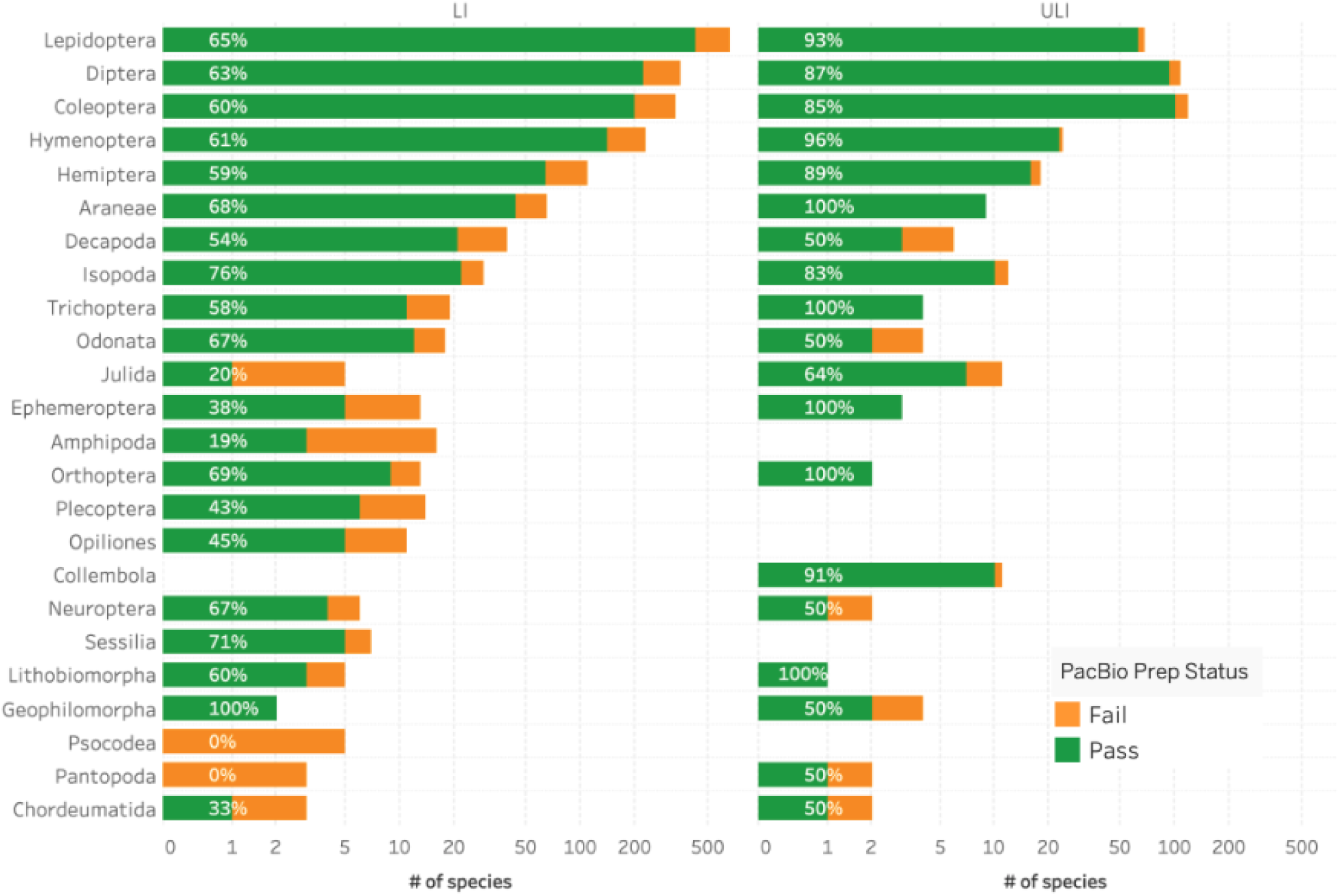
Arthropod fragmentation success by order. The bar chart summarises the DNA fragmentation success per species across Arthropod orders, subjected via the LI (left) or ULI (right) submission types. Species progressed under both submission types are included in both bars. The results represent the best DNA fragmentation outcome per species and submission type, determined using the hierarchy: Pass > Fail. The number inside each bar represents the percentage of species that have passed extraction within the orders. To account for the wide range in species counts, a logarithmic scale is used, and orders with fewer than five species are excluded from this visualisation but are available in the supplementary material (Figure S2).

#### Plants

Plant samples are typically fresh leaf material that has been collected in relative abundance, supported by the use of 7.6 mL tubes. Tissue availability is usually not a limiting factor for plants, other than particular taxonomic groups such as the Bryophytes.

We use the ‘Plant MagAttract’ [29] protocol routinely for DNA extraction protocol from all plant species (Figure 1). It is efficient at extracting HMW DNA from a wide range of species to an extent adequate for long-read sequencing. Plant samples that fail to produce sequenceable HMW DNA from the Plant MagAttract extraction protocol are processed through the Plant Organic Extraction (POE) protocol [30]. Species extracted with the Plant MagAttract v.4 protocol can result in a poor DNA profile that is significantly improved when the same species is extracted with the POE protocol (Figure 10). The POE protocol is mid-throughput and requires more time and expertise in the laboratory, and for this reason it is employed only as a second-measure attempt for recalcitrant species. Work is underway to identify prior to extraction which species would most benefit from proceeding directly to the POE extraction method.

**Figure 10.**
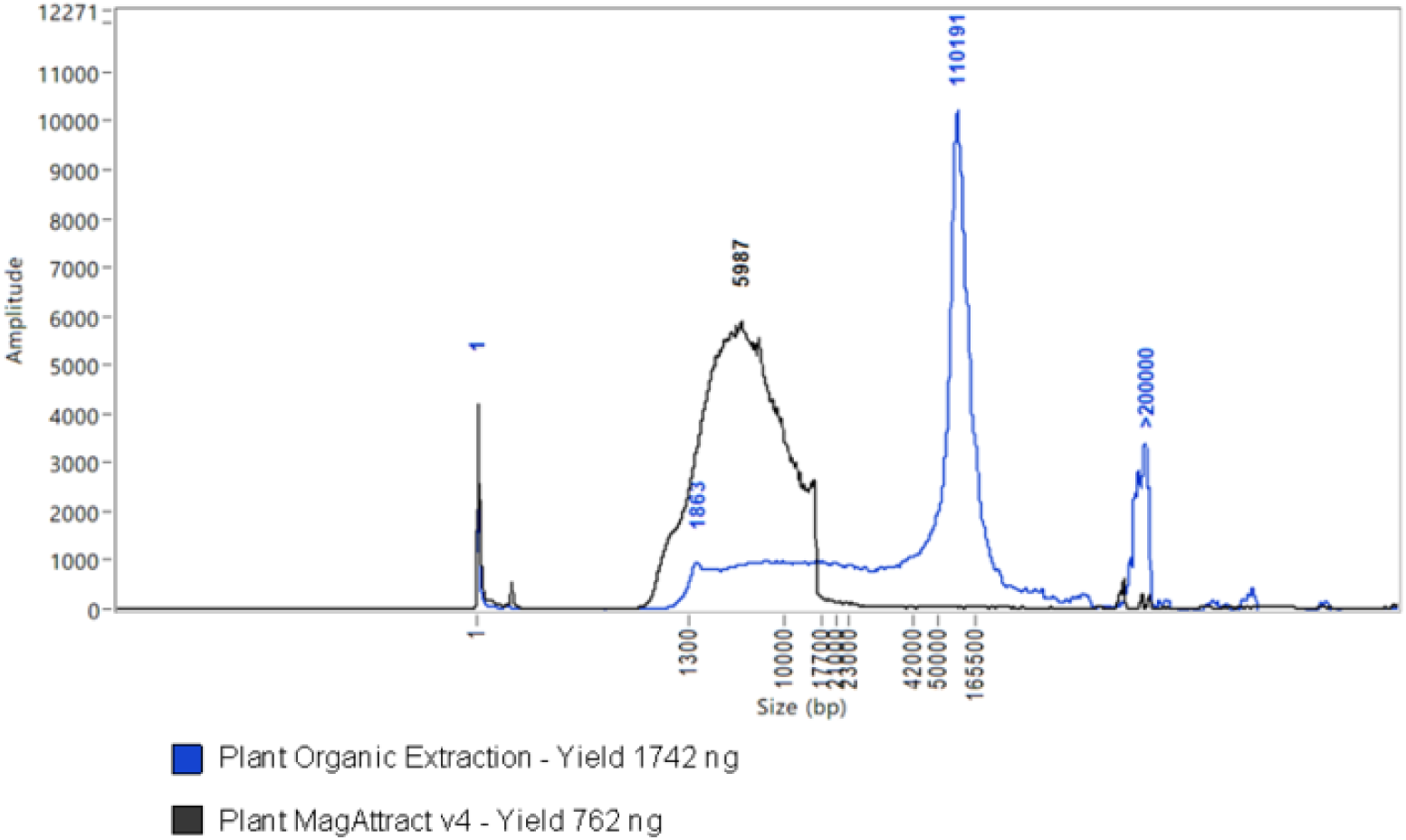
Overlaid FemtoPulse molecular weight profiles of two DNA extractions from *Thymus drucei* (Lamiales; wild thyme) with two different DNA extraction protocols. The trace from the Plant MagAttract extract from 67 mg tissue shows primarily a wide LMW peak at 5.8 kb, whereas the POE protocol extract from 65 mg tissue shows a strong peak at 110 kb with slight smear down to 1.8 kb and further strong peak in the ultra HMW zone (>200 kb).

We have processed 998 species covering 63 orders, 166 families and 564 genera within the Plant taxonomic group, including 927 vascular (Tracheophyta and Streptophyta) and 71 non-vascular (Bryophyta and Marchantiophyta) species. High success rates have been observed across the plant orders with 91% of all species extracted having passed through to subsequent processes. However, many of the successfully extracted plant species have failed at later stages including fragmentation, library preparation, or sequencing. Plant species that have failed twice at any of these points have been selected and processed through the POE protocol. The switch to use of the POE protocol is clearly of benefit to some groups, such as the Saxifragales, where the pass rate changes from 56% with MagAttract to 100% with POE (Figure 11).

**Figure 11.**
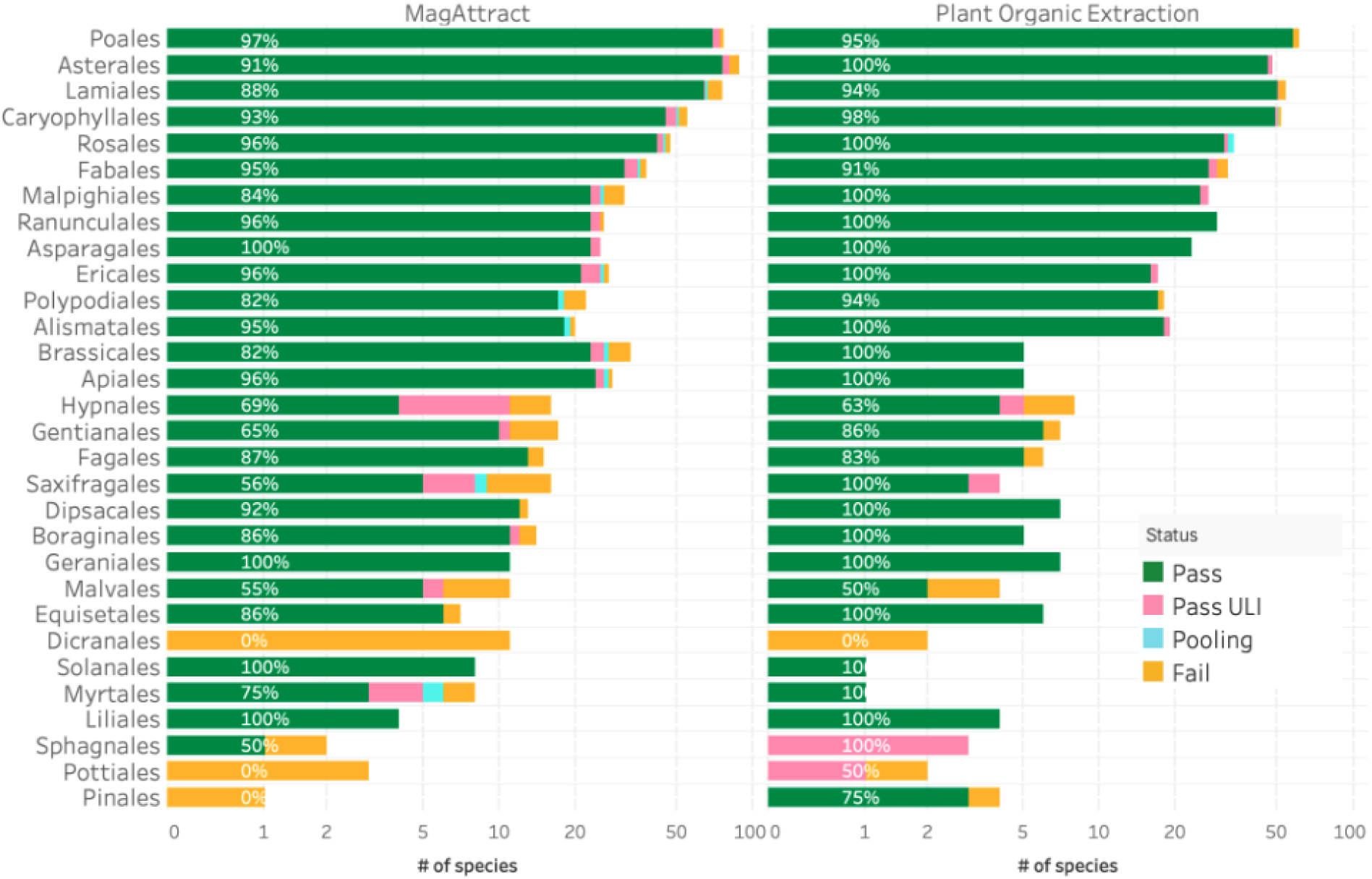
Plant MagAttract and Plant Organic Extraction (POE) DNA extraction success metrics by order. The bar chart summarises the DNA extraction success per species across Plant orders, extracted via MagAttract or Plant Organic Extraction protocols. Species extracted with both protocols are included in both bars. Species extracted with protocols other than Plant MagAttract or Plant Organic Extraction are not represented. The results are categorised as: Pass – DNA sufficient for sequencing achieved; Pass ULI - DNA sufficient for sequencing with ultra low input achieved; Pooling - two DNA extractions were pooled to meet QC threshold; Fail - extractions have failed to provide sufficient DNA to proceed. The results represent the best DNA extraction outcome per species and extraction protocol, determined using the hierarchy: Pass > Pass ULI > Pooling > Fail. The number inside each bar represents the percentage of species that have passed extraction within the orders, including Pass, Pooling and Pass ULI categories. To account for the wide range in species counts, a logarithmic scale is used, and orders with fewer than five species are excluded from this visualisation but are available in the supplementary material (Figure S3).

Fragmentation of DNA extracted from plant material is usually not problematic, with a success rate of 82%, calculated using unique species regardless of the submission protocol. For species for which both protocols have been used, the LI and ULI submission protocols have had a success rate of 80% and 93% respectively (calculated from data in [4]).

Bryophytes have been challenging due to their low tissue availability as an individual is often <15 mg, whereas the usual input for plant DNA extraction methods is 50 mg. Modification of protocols to minimise tissue loss and maximise DNA recovery have been successful, coupled with the ULI library prep method as the genome sizes are often <1 Gb.

A number of plant species remain challenging, with neither MagAttract nor POE DNA extraction protocols providing the required DNA yield or quality for long read sequencing. A pre-lysis hypertonic sorbitol wash has been developed [31] to remove interfering chemical contaminants present within the cytosol of plant specimens prior to lysis. Sorbitol is an osmotically active sugar alcohol capable of ‘drawing out’ the cytosol of plant tissues homogenates without interrupting the nuclear membrane. When a sorbitol wash is successful, a previously recalcitrant sample’s lysate should be absent of both viscosity, browning or other unfavourable characteristics. Initial results have shown that this protocol can significantly improve the quality of DNA extractions, and the CCS yield.

#### Fungi (including Lichens)

Most fungal samples received had been cultured from samples collected in the field. The samples arrived as cell pellets with low tissue mass and size presenting challenges for DNA extraction. Mycelium samples can also be challenging due to low density of nuclei in the tissue. DNA extractions for these fungi have been of consistently low yield, and often of low quality in terms of fragment length resulting in a high ‘Fail’ rate (Figure 12). On occasions that high yields have been achieved, the sequencing of these samples has been poor. Given the typically small genome size of fungi (typically ∼40 Mb) and the relatively low yields achieved, the ULI library prep method has been the standard option for fungi samples. The ideal amount of DNA to start this process is 100 ng, although samples with >25 ng of DNA within the ULI fragment range are progressed. The optimised automated Plant Magattract protocol (v.4) [29] has provided increased DNA yield and improved FemtoPulse profile and is therefore now the protocol of choice. Post-extraction, the majority of fungi samples are directed toward the g-TUBE fragmentation method [26], this is an efficient and effective process resulting in a high pass rate for unique fungi species of 79% (calculated from data in [4]). The amplification in the ULI library generation process is being utilised here to aid sequencing, rather than accounting for a very low DNA input amount, as native fungal DNA often produces poor sequencing yields. To optimise the amplification process for this purpose, we are currently exploring a reduction in the number of PCR cycles and trialling different enzymes.

**Figure 12.**
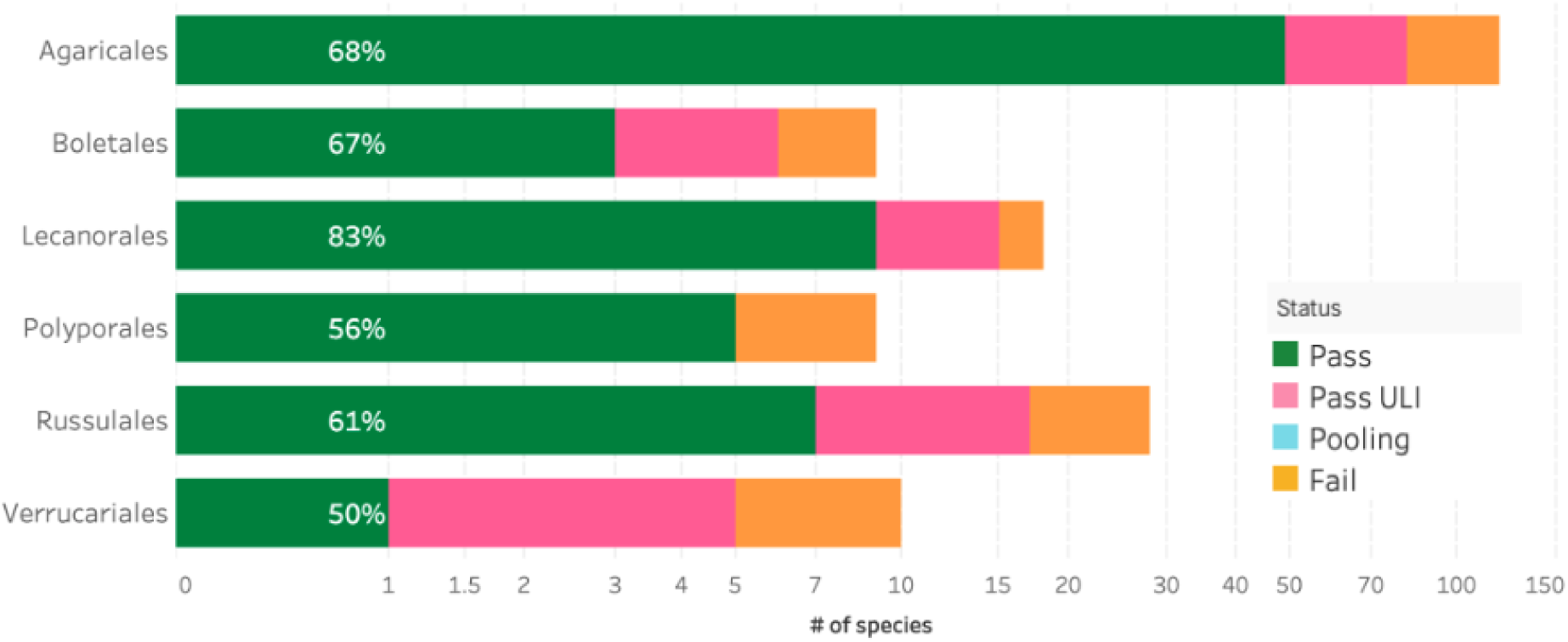
Fungi DNA Extraction success metrics by order. The bar chart summarises the DNA extraction success per species across Fungi orders. The results are categorised as: Pass – DNA sufficient for sequencing achieved; Pass ULI - DNA sufficient for sequencing with ultra low input achieved; Pooling - two DNA extractions were pooled to meet QC threshold; Fail - extractions have failed to provide sufficient DNA to proceed. The results represent the best DNA extraction outcome per species, determined using the hierarchy: Pass > Pass ULI > Pooling > Fail. The number inside each bar represents the percentage of species that have passed extraction in any way, not those that failed. To account for the wide range in species counts, a logarithmic scale is used, and orders with fewer than five species are excluded from this visualisation but are available in the supplementary material (Figure S4).

#### Chordates

The routine processing of chordates is highly efficient, resulting in a 96% pass rate for species at DNA extraction (Figure 6). The DNA extraction status for orders within the group reveals that this success is general, with no clear trends (Figure 13). Chordates were processed at 89% pass rate (calculated from data in [4]) at DNA fragmentation, when results for both protocols (g-TUBE and Megaruptor) are combined at the species level.

**Figure 13.**
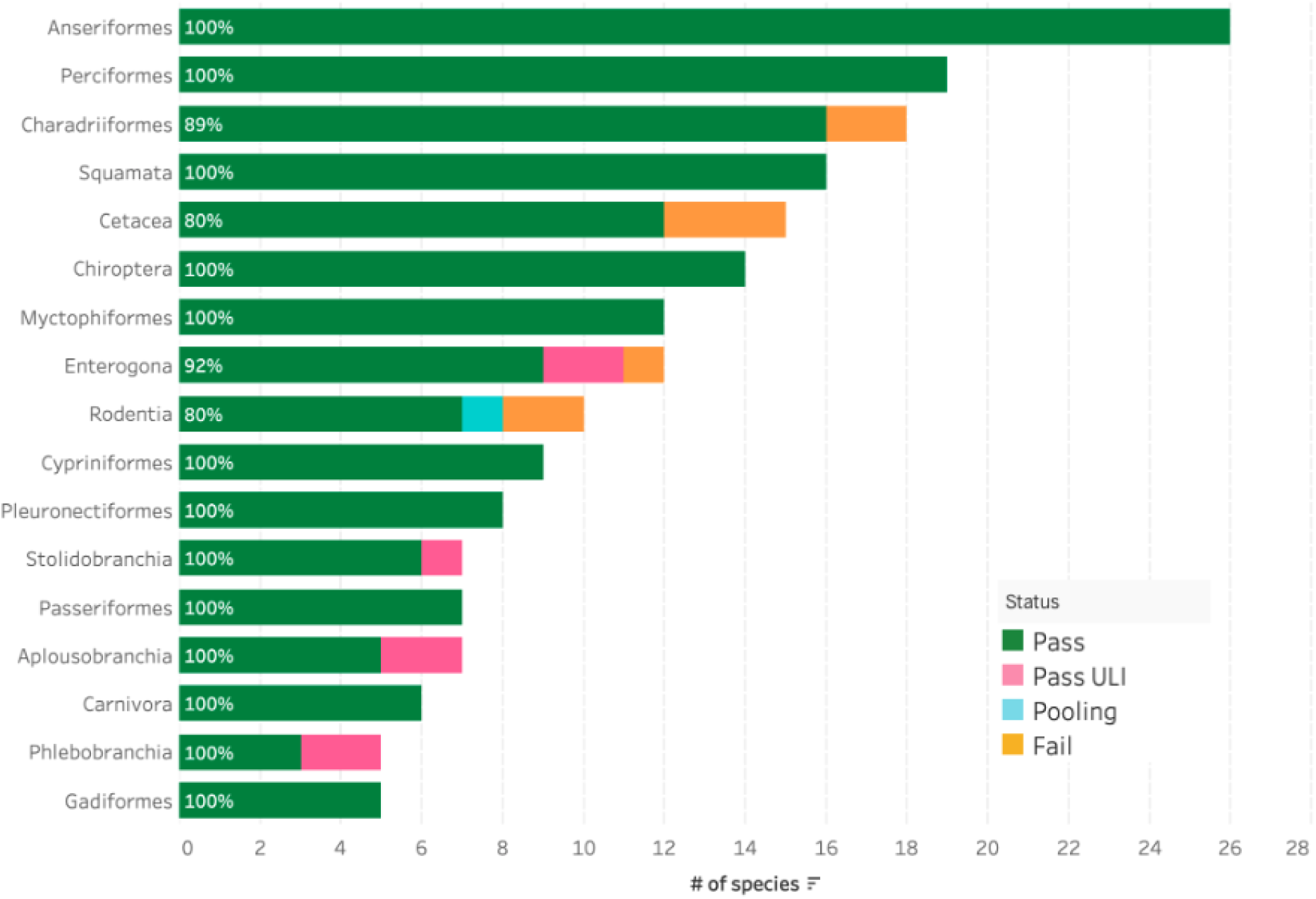
Chordate DNA Extraction success metrics by order. The bar chart summarises the DNA extraction success per species across Chordata orders. The results are categorised as: Pass – DNA sufficient for sequencing achieved; Pass ULI - DNA sufficient for sequencing with ultra low input achieved; Pooling - two DNA extractions were pooled to meet QC threshold; Fail - extractions have failed to provide sufficient DNA to proceed. The results represent the best DNA extraction outcome per species, determined using the hierarchy: Pass > Pass ULI > Pooling > Fail. The number inside each bar represents the percentage of species that have passed extraction in any way, not those that failed. To account for the wide range in species counts, orders with fewer than five species are excluded from this visualisation but are available in the supplementary material (Figure S5).

The collection of chordate samples is legally and ethically challenging, and due to this, a significant number of samples from chordate species are provided from specimens that have been found dead, or small tissues collected from live individuals. The use of preservative solutions in lieu of snap-freezing is common for chordate samples. To ensure proper fixation it is recommended that a 1:10 ratio of tissue:fixative volumes is used. Ear punches and punch biopsies are a very useful form of non-lethal sample collection for chordate species; though not the most successful for DNA extraction, disruption via the cryoPREP helped maximise DNA yield and quality. Fish and bird blood are amongst the best performing tissues for HMW DNA extraction and are processed using the Nanobind HMW DNA extraction - nucleated blood protocol [32]. This manual protocol requires inputs ranging from 5-25 µl of nucleated blood, flash frozen or stored in ethanol at −80°C, from birds, fish or amphibians, and yields around 10-40 µg of HMW DNA. An automated version of this protocol [33] permits high throughput extraction of nucleated blood samples. Samples collected at necropsy often contain only degraded DNA. In these situations, it is possible to extract and then perform a stringent 0.45X SPRI to remove any remaining RNA or LMW DNA, and progress directly to library preparation and sequencing, bypassing shearing, if the fragment size profile is already degraded.

#### Protists

This taxonomic group poses a unique challenge due to the diversity of the species it contains, from microalgae to dinoflagellates, the nature of cell walls and exoskeletons, and the relative size of both individuals and genomes [34].

Protist samples have typically been provided as cell pellets from cultured strains. Because of the diversity of culture conditions required by different species this results in pellets with a wide range in mass and cell number per mg weight. This diversity makes it hard to standardise input amounts for DNA extraction. Although not yet fully optimised, the currently preferred process begins with a cell pellet of 50 mg, disrupted with the cryoPREP. DNA is extracted using the Plant MagAttract v4 extraction protocol [29,35]. The use of the cryoPREP provides increased yields compared to power mashed samples, and the adoption of Plant MagAttract v4 lysis also increases yield due to the better lysis of cell wall structures present in many protists and microalgae. These measures have resulted in an overall success rate in extraction for protists of 83% (Figure 6) with an uneven distribution between orders (Figure 14).

**Figure 14.**
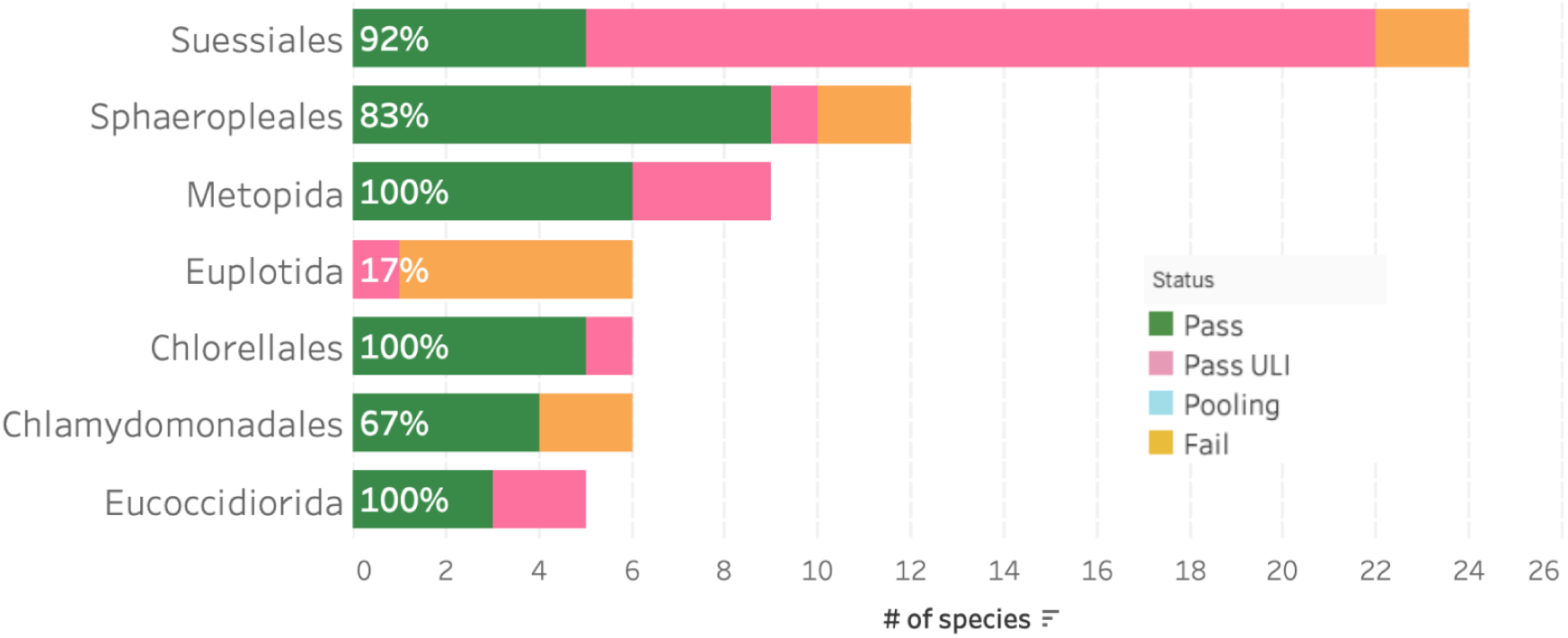
Protist DNA extraction success metrics by order. The bar chart summarises the DNA extraction success per species across Protist orders. The results are categorised as: Pass – DNA sufficient for sequencing achieved; Pass ULI - DNA sufficient for sequencing with ultra low input achieved; Pooling - two DNA extractions were pooled to meet QC threshold; Fail - extractions have failed to provide sufficient DNA to proceed. The results represent the best DNA extraction outcome per species, determined using the hierarchy: Pass > Pass ULI > Pooling > Fail. The number inside each bar represents the percentage of species that have passed extraction in any way, not those that failed. To account for the wide range in species counts, orders with fewer than five species are excluded from this visualisation but are available in the supplementary material (Figure S6).

When species are identified as being cultured in axenic or low bacteria conditions by sample providers and with a predicted genome size of below 1 Gb we are generally able to assemble genomes with ULI sequencing and Hi-C data. Protist samples predominantly progress toward the ULI route, achieving a success rate of 83%. In contrast, we have lower success rates (57%) with LI shearing protocols (calculated from data in [4]). Overall, our combined success rate is 74%, with future work aiming towards improvement of the ULI pipeline. The unusual genome structure of some protist species, for example ciliates, provide an extra challenge for fragmentation and size selection. Chromosomes are present in the range of 5 kb to 20 kb and these would be removed using current size selection protocols, future methods to efficiently sequence these fragments may include fractionation of DNA extracts.

Many protists feed on bacteria, or require their presence for growth, and for this reason samples can be a mixture of protist and bacterial cells in culture. Sequencing yields for protist ULI samples may be very good, achieving over 24 Gb per Sequel IIe cell. However, up to 99% of these reads can originate from co-cultivated bacteria within the sample rather than the target protist. The importance of working with collectors to reduce this bacterial load is therefore fundamental to the progression of protist samples.

Future work in this area will focus on assessment of dual extraction protocols, aiming to extract easily lysed organisms and remove this DNA in a first pass, followed by a stronger chemical or physical cell lysis and DNA extraction for the remaining sample.

#### Other Metazoa and Macroalgae

The paraphyletic group of “other metazoa and macroalgae” contains a multitude of different phyla, predominantly a mix of marine and terrestrial invertebrates, but also including macroalgae. This grouping is largely based on the focus of the species collectors, and their access to species whilst sampling in marine environments. The large diversity of the species within this polyphyletic grouping provides many challenges and opportunities for new developments.

Samples within this group are homogenised using either cryoPREP or powermashing, based on the weight of the tissue available as described in the standard guidelines [16] and then subjected to the automated MagAttract extraction process [24]. Species are matched with ideal extraction protocols with an overall pass rate of 79% (Figure 6). This is relatively low compared with other taxonomic groups and is not spread evenly, with for example Mollusca showing a high success rate of 87%, whereas Platyhelminthes only achieve 29% (Figure 15).

**Figure 15.**
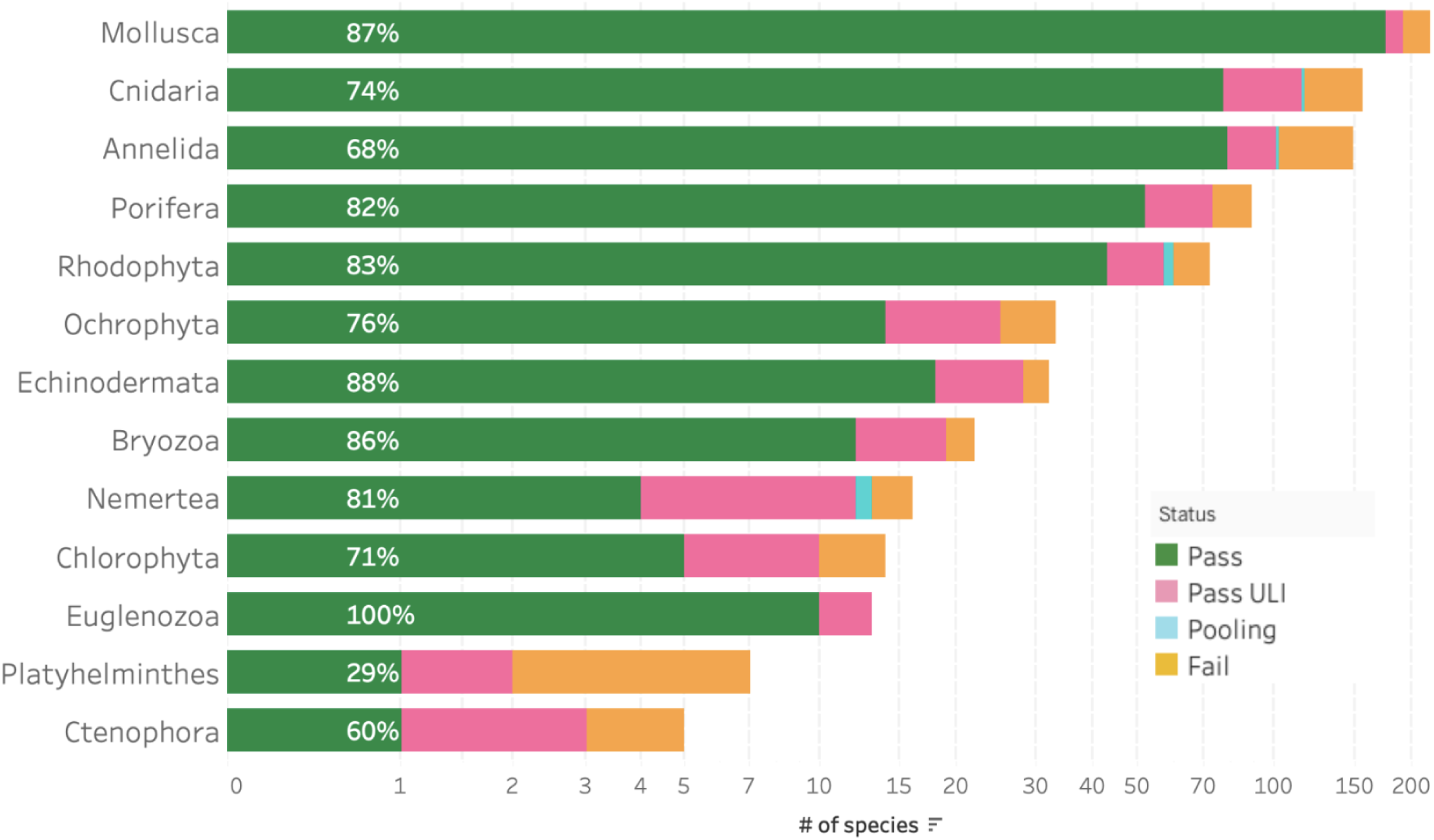
Other metazoa and macroalgae extraction success metrics by taxon group. The bar chart summarises the DNA extraction success per species across other metazoa and macroalgae taxon groups. The results are categorised as: Pass – DNA sufficient for sequencing achieved; Pass ULI - DNA sufficient for sequencing with ultra low input achieved; Pooling - two DNA extractions were pooled to meet QC threshold; Fail - extractions have failed to provide sufficient DNA to proceed. The results represent the best DNA extraction outcome per species, determined using the hierarchy: Pass > Pass ULI > Pooling > Fail. The number inside each bar represents the percentage of species that have passed extraction in any way, not those that failed. To account for the wide range in species counts, a logarithmic scale is used, and taxon groups with fewer than five species are excluded from this visualisation but are available in the supplementary material (Figure S7).

The results of fragmentation processes for the orders contained in other metazoa show a pass rate of 74% (calculated from data in [4]). ULI is a useful option when fragmentation results are poor as many of the fragmentation failures are associated with a low yield or a poor profile. ULI libraries dominate for the majority of orders, particularly Cnidaria where 54 species have been processed via LI and 108 via ULI library prep. Mollusca are the exception for this trend, with 186 species processed for LI and only 66 for ULI.

#### Molluscs

While many molluscs pass *via* the routine MagAttract protocol [20,24], the Nanobind [36] method is used as the second option for those that fail (Figure 1). An example of this is *Colus gracilis* (Gastropoda; Graceful whelk), which failed consistently for DNA quality when extracted using the MagAttract Protocol [24] (Figure 16). However, when processed using the Nanobind protocol [36] DNA with a high molecular weight peak and a profile suitable for sequencing was obtained. The Nanobind extraction also increased the overall yield tenfold. This result may be conflated by a difference in the tissue preparation for these protocols, as the MagAttract samples were disrupted in the cryoPREP (Covaris, Woburn, MA) whilst the Nanobind samples were finely diced with a scalpel as per the protocol.

**Figure 16.**
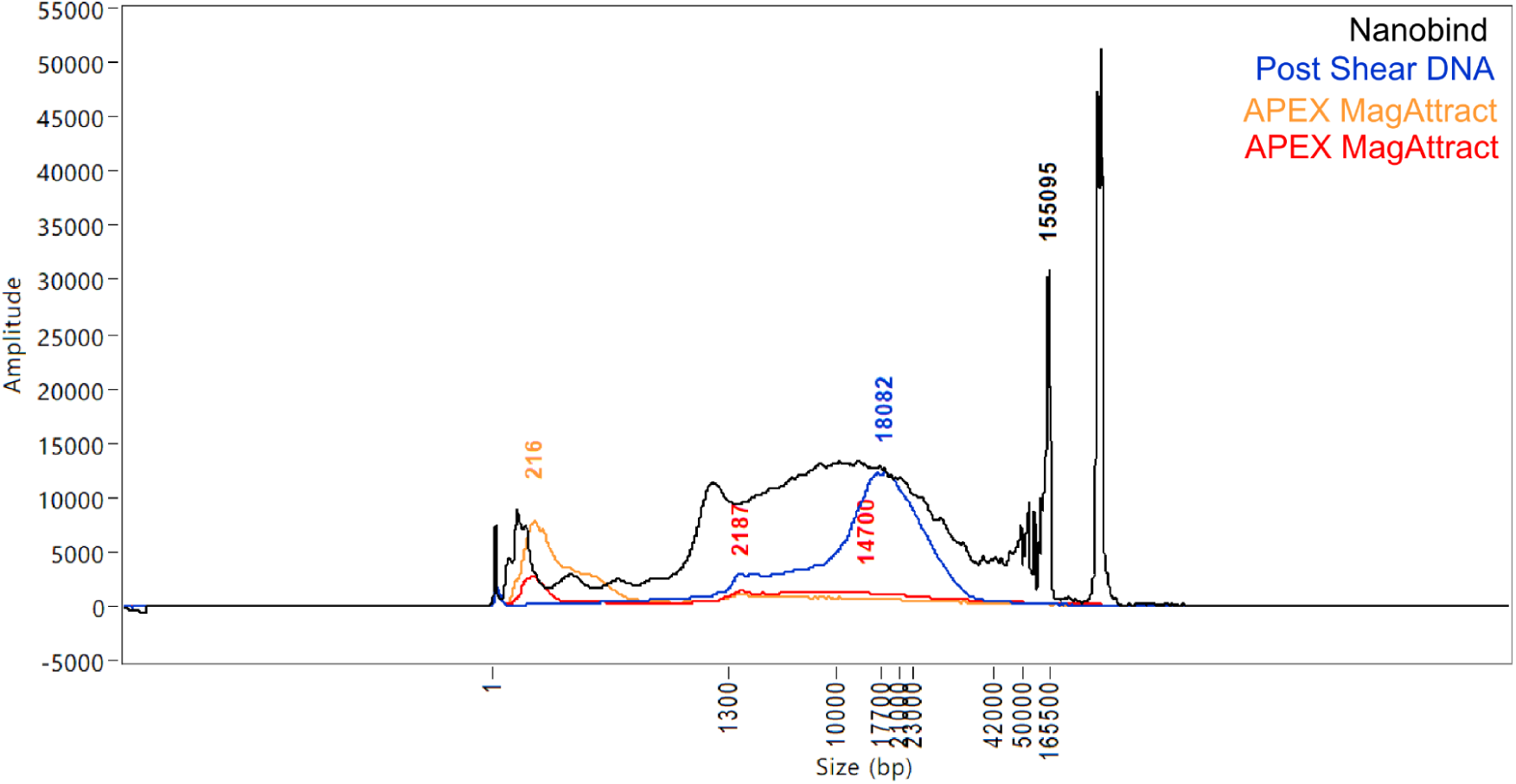
FemtoPulse profiles of the mollusc *Colus gracilis* (Gastropoda) DNA extracts following different protocols. The samples extracted using the automated MagAttract protocol (Yellow and Red) yielded only LMW DNA and are not suitable for progression. The results from the Nanobind protocol (Black) show a significant improvement, both in the abundance of HMW DNA and also the absence of LMW DNA. After fragmentation and clean up of the Nanobind extracted DNA, the resulting peak fragment size of 18 kb (Blue) was ideal for progression to library prep.

#### Cnidaria

Cnidaria samples have also proved challenging, with corals causing difficulties during sample homogenisation, and jellyfish yielding low quality and quantity DNA following routine DNA extraction. One of the big challenges of extracting DNA from corals has been in disrupting hard stony corals into a fine powder that facilitates extraction. The deployment of the Fast-prep96™ has enabled faster and more complete disruption of coral tissue *via* a scalable method using 4 ml polycarbonate vials and a single 6mm zirconium oxide grinding bead, and otherwise following the plant bead beating protocol [19]. This approach to sample disruption has improved the MagAttract extraction success of hard corals, resulting in an increase in samples that yielded DNA suitable for LI or ULI sequencing. The bead-beaten coral samples have also been successfully used for Hi-C cross linking and subsequent library preparation, completing the data set required for reference level genome assembly.

For salps and jellyfish, a new extraction protocol was developed, using the recommended lysis steps of the Omega Bio-Tek E.Z.N.A. Mollusc and Insect DNA kit (Item: D3373-00S from Omega Bio-Tek, Norcross, GA.) combined with the SpeedBead-based extraction method used in the POE protocol, which generated a higher quality and quantity of DNA. This protocol required inputs of 100 - 200 mg of fresh frozen tissue; lower input amounts and ethanol preserved tissues could also be used, however the resulting DNA yield may be lower. The DNA extracted using the Modified Omega Bio-Tek E.Z.N.A. protocol [37] was suitable for either ULI or LI sequencing, enabling the reference level assembly of multiple salp and jellyfish genomes.

#### Porifera

Initially, processing through the routine protocols of cryoPREP [18] and MagAttract v2 [38] yielded a significant portion of LMW DNA within the extract. Research into homogenisation methods that have been used identified the “squeeze” method [27], which aims to maintain the integrity of the sponge cells whilst removing them from their skeletons (siliceous and calcareous spicules embedded in collagenous protein matrices). Samples of *Eunapius fragilis* (a freshwater demosponge) extracted with and without “squeezing” showed significantly increased yields of high molecular weight DNA in the squeezed sample. The cells separated via the squeeze method have also been successfully used to generate Hi-C data, so all Porifera are now processed using the squeeze method.

#### Macroalgae

Initial work with macroalgal samples (Chlorophyta, Ochrophyta and Rhodophyta) began with tissue disruption via the cryoPREP [18] followed by the POE protocol [29], yielding DNA that was of both poor quality and quantity. The samples were characterised by their tendency to become very viscous upon cell lysis, forming a gel like substance in the tube during extraction which significantly hindered further processing. This is due to the large polysaccharide content of the algae. The typical approach for macroalgae with a genome size <1 Gb is therefore the ULI library prep method after the POE extraction method with a lower tissue input of 25mg in order to reduce the amount of contaminants in the sample.

## Conclusion

The Sanger Tree of Life programme has scaled reference genome assembly production and has released over 2000 chromosomally-resolved genome reference assemblies as of February 2025. We aim to further increase genome production year on year, and standardisation, refinement and streamlining of laboratory processes has been fundamental for our continual improvements. The homogenisation methods, extraction protocols, and shearing processes discussed here are enabling genome assemblies from a great diversity of species. In addition to release of the data freeze used to produce the summary statistics presented here [4], in order to further assist others working in the field, we have also made the raw data from the Tree of Life laboratory work available via a searchable online ‘Portal’ at links.tol.sanger.ac.uk/datasets/tol-lab-data. This link is continuously updated with the work underway and thus provides access to laboratory information as soon as the work has been completed. We hope this will be useful to examine both the details behind the summaries offered in this paper but also to explore the protocols used on future samples. For example, if a researcher is working on a challenging species that is a close relative of a species that has come through the Tree of Life, the portal could be explored to understand which HMW DNA extraction protocols worked or did not work and thus save time in testing a variety of approaches. Alternatively, where a researcher has access to multiple tissue types for work, the Portal may provide information as to how related species and tissue types have performed in extraction and downstream sequencing, informing decision making.

Our experience shows that building high quality reference genome assemblies is achievable for the majority of species that have been collected alive and preserved using best practice (snap freezing in most cases), and have a suitable tissue availability-to-genome size ratio. Challenges remain in certain taxonomic areas, especially for species with large genomes and small body sizes. The new Ampli-Fi option from PacBio requires only 1 ng of sheared DNA to provide data for up to 3 Gb genome size and may help overcome some of these challenges. Best practice is to avoid amplification whenever possible, and even here, the requirements for input DNA amounts are regularly decreasing with a recent four-fold decrease in the amount of DNA needed for LI PacBio. The Sanger Tree of Life core lab biobanks all DNA aliquots including those that did not meet the quality and yield required to progress to sequencing at the time of their extraction. With these recent advances, we will now return to biobanked DNA extracts that previously did not meet required yields as sequencing these to sufficient coverage may now be achievable.

For species with picogram level DNA content, phi29 replicase amplification can be used on single meiofaunal organisms to generate long-insert library DNA. Picogram input Multimodal Sequencing (PiMmS) [23] delivers a PacBio or ONT long-read compatible amplified DNA sample and full length cDNA from a single specimen. This has proven successful for a number of species [39], and work will continue to standardise and ramp up the use of this type of method.

Future work will continue to focus on the species that fail at different stages of the process, to develop and implement methods for the processing of smaller samples with as little amplification as possible, and to explore the merits of different sequencing technologies. We will continue to share our protocols and findings as soon as possible in the hope that global biodiversity genomics efforts might benefit.

## ACKNOWLEDGEMENTS

All authors as well as the laboratory work discussed above were funded by the Wellcome Sanger Institute Quinquennial Review award 2021-2026 to the Wellcome Sanger Institute (220540/Z/20/A). In addition, the majority of genome production for species among the first 2000 discussed here was supported by Wellcome through the Darwin Tree of Life Discretionary Award (218328) and by the Gordon and Betty Moore Foundation through the Aquatic Symbiosis Genomics Project (Grant ID: GBMF8897, https://doi.org/10.37807/GBMF8897).

We thank the many hundreds of people who collected and identified species on behalf of the Darwin Tree of Life and Aquatic Symbiosis Genomics Projects, and the many colleagues in these projects who shared their best methods with us. We also thank the staff of the Wellcome Sanger Institute Scientific Operations teams who contributed to extractions, and conducted library preparation and sequencing.

