## Supplementary material for "On the path to reference genomes for all biodiversity: lessons learned and laboratory protocols created in the Sanger Tree of Life core laboratory over the first 2000 species": Figure S1

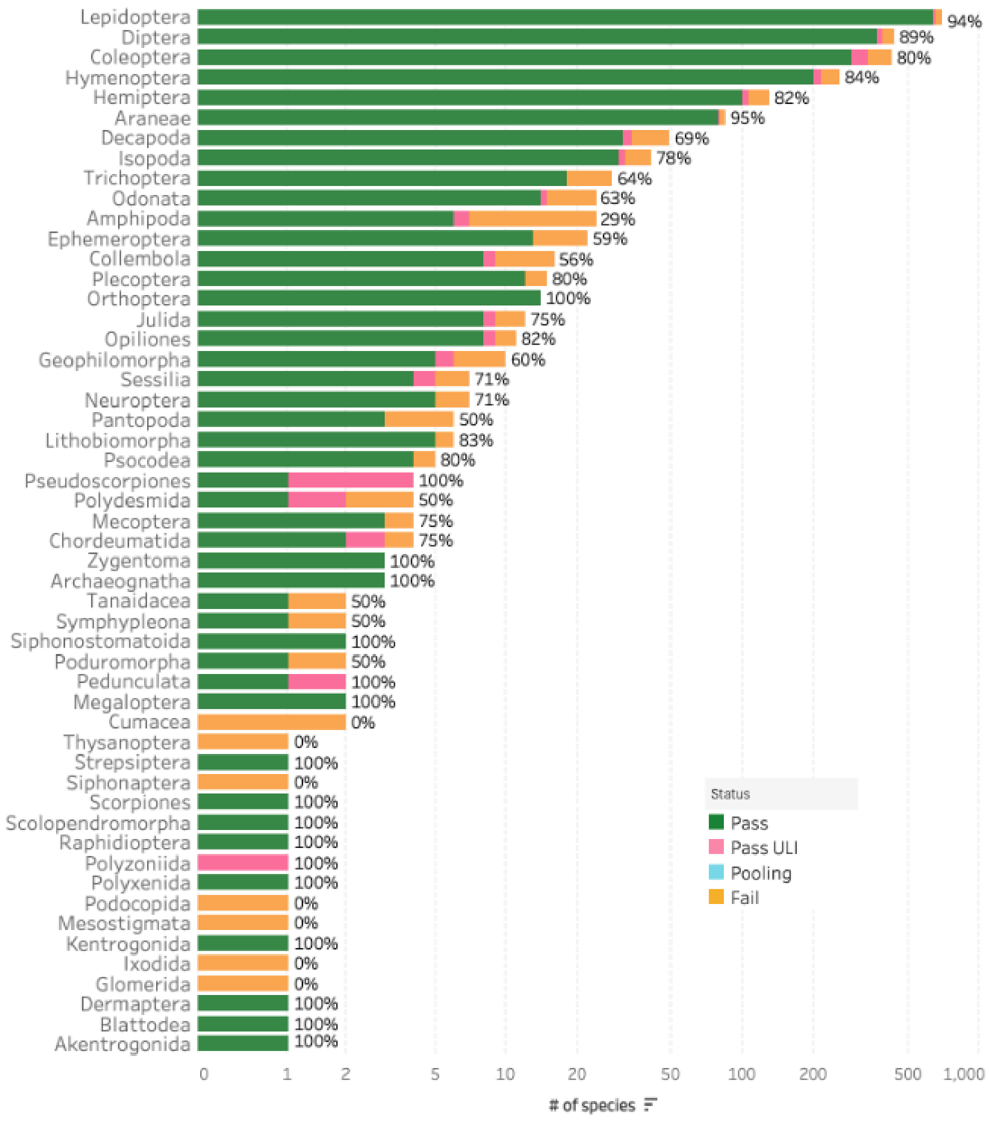


**Figure S1. Arthropod DNA extraction success metrics by order**

The bar chart summarises the DNA extraction success per species across Arthropod orders. The results are categorised as: Pass – DNA sufficient for sequencing achieved; Pass ULI - DNA sufficient for sequencing with ultra low input achieved; Pooling - two DNA extractions were pooled to meet QC threshold; Fail - extractions have failed to provide sufficient DNA to proceed. The results represent the best DNA extraction outcome per species, determined using the hierarchy: Pass > Pass ULI > Pooling > Fail. The number to the right of each bar represents the percentage of species that have passed extraction within the orders, including Pass, Pooling and Pass ULI categories. To account for the wide range in species counts, a logarithmic scale is used.
