## Supplementary material for "On the path to reference genomes for all biodiversity: lessons learned and laboratory protocols created in the Sanger Tree of Life core laboratory over the first 2000 species": Figure S2

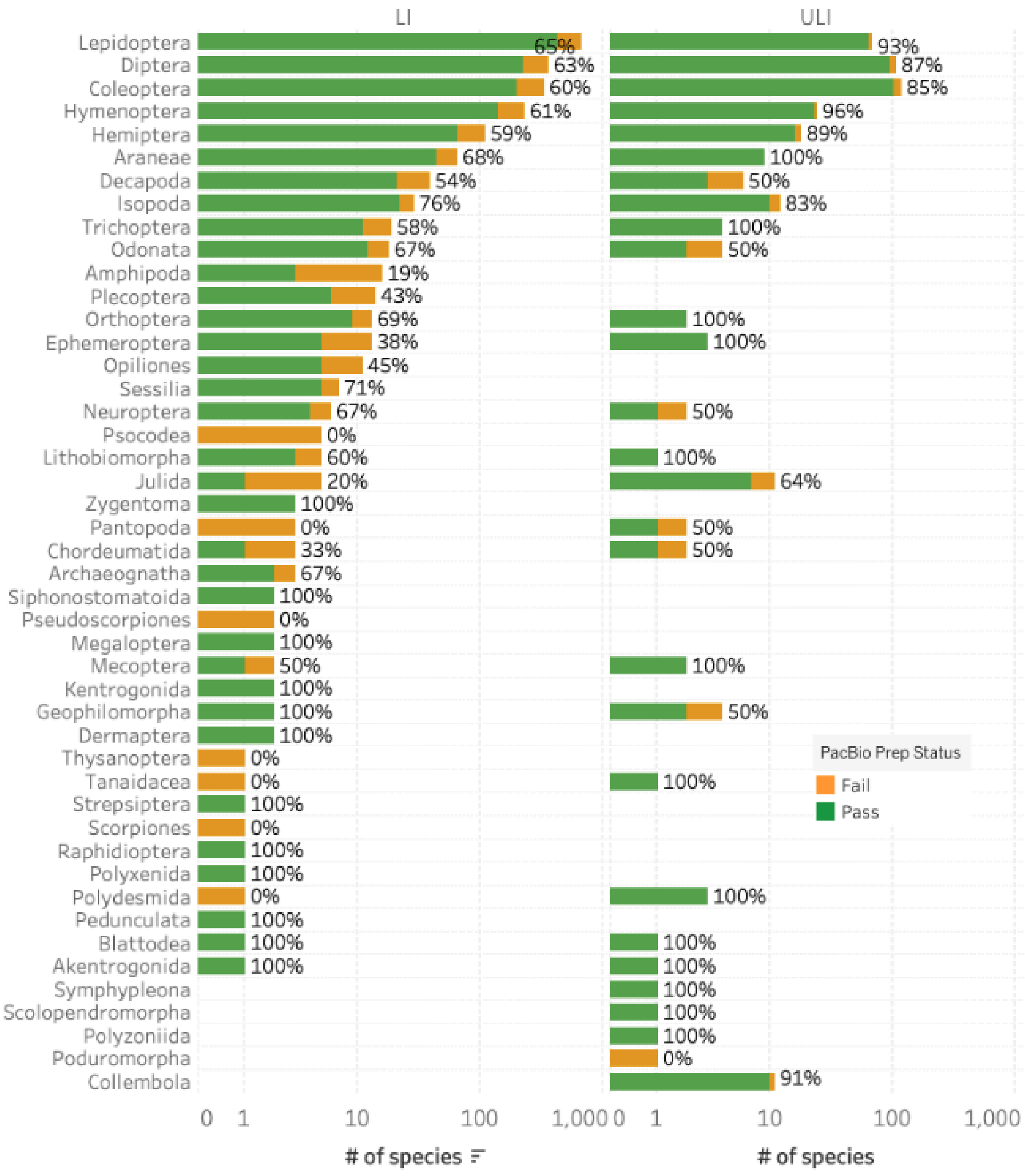


**Figure S2. Arthropod fragmentation success by order**

The bar chart summarises the DNA fragmentation success per species across Arthropod orders, subjected via the LI or ULI submission types. Species progressed under both submission types are included in both bars. The results represent the best DNA fragmentation outcome per species and submission type, determined using the hierarchy: Pass > Fail. The number to the right of each bar represents the percentage of species that have passed extraction within the orders. To account for the wide range in species counts, a logarithmic scale is used.
